# A perturbation proteomics-based foundation model for virtual cell construction

**DOI:** 10.1101/2025.02.07.637070

**Authors:** Rui Sun, Liujia Qian, Yongge Li, Honghan Cheng, Zhangzhi Xue, Xuedong Zhang, Lingling Tan, Yuecheng Zhan, Wenbin Hu, Qi Xiao, Zhiwei Liu, Guangmei Zhang, E Weinan, Peijie Zhou, Han Wen, Yi Zhu, Tiannan Guo

## Abstract

Building a virtual cell requires comprehensive understanding of protein network dynamics of a cell which necessitates large-scale perturbation proteome data and intelligent computational models learned from the proteome data corpus. Here, we generate a large-scale dataset of over 38 million perturbed protein measurements in breast cancer cell lines and develop a neural ordinary differential equation-based foundation model, namely ProteinTalks. During pretraining, ProteinTalks gains a fundamental understanding of cellular protein network dynamics. Our model encodes protein networks and exhibits consistently improved predictive accuracy across various downstream tasks, highlighting its generalization capabilities and adaptability. In cancer cells, ProteinTalks robustly predicts drug efficacy and synergy, identifies novel drug combinations, and, through its interpretability, uncovers resistance-associated proteins. When applied to more complex system, patient-derived tumor xenografts, ProteinTalks predicts potential responses to drugs. Its integration with clinical patient data enhances the prognosis prediction of breast cancer patients. Collectively, we present a foundational model based on proteome dynamics, offering potential for various downstream applications, including drug discovery, and providing a basis for developing virtual cells.

## Introduction

Virtual cell refers to a computational model of a physical cell, designed to simulate and predict cellular processes *in silico* ^1-3^. Constructing a virtual cell requires thorough time-resolved measurements of all cell ingredients, reflecting the cellular functions and dynamic behaviors ^4^. Protein networks are essential for understanding cell biology and disease mechanisms, providing insights into disease progression and informing treatment strategies ^5,6^. However, large-scale proteomic data, particularly including dynamic information, remains extremely sparse compared to transcriptomic data. In drug discovery, targeting specific proteins without considering their network context may result in limited efficacy or unintended side effects ^7^. Consequently, systematic analysis and modeling of proteomic dynamics are crucial for identifying novel drug targets, designing more effective and precise therapies, and ultimately developing comprehensive virtual cell models.

Perturbation proteomics provides a powerful approach to decipher complex protein network dynamics, aiding drug discovery by uncovering drug mechanisms of action (MOAs) ^8-12^. With advances in high-throughput proteomics ^13^ enabled by data-independent acquisition mass spectrometry (DIA-MS) ^14^, generating large-scale perturbation proteomics datasets is now achievable ^9^, paved way for utilizing large scale pre-training techniques to describe the protein dynamical space.

Here, we integrated a multidimensional perturbation approach to comprehend the complexity of interconnected drug-protein systems. Utilizing a 96-well-based high-throughput platform, we perturbed breast cancer cell lines with clinically relevant drugs and their combinations, and subsequently obtained over 38 million perturbed protein measurements, as well as cell morphological and viability data. Based on this large-scale proteome data corpus, we further developed a dynamical foundation model called ProteinTalks, based on ordinary differential equation (ODE) network models ^15^, to predict perturbed proteomic networks, then extend to identification of essential proteins associated with drug efficacy, and characterization of cellular response after drug treatment. Since the dynamical information was explicitly integrated in the model architecture, our model demonstrated excellent generalizability and adaptability in multiple downstream applications in cell lines, mouse models and patients, with significantly fewer parameters. Overall, this study introduces a paradigm shift to combine large scale perturbation data and dynamical neural network to expand the concept of foundation models beyond scale to include the dimension of time. We believe such rationale will serve as an important alternative to effectively make use of limited measurements to build limitless virtual cell models across space and time.

## Results

### Generation of perturbation proteome datasets for building a foundation model

As a proof of concept, and to be directly related to drug development, we focused specifically on cancer cell lines, specifically breast cancer cells (**Figure 1A**). We selected 16 commonly used TNBC cell lines and two non-TNBC cell lines (**Table S1A**). We also collected 63 FDA-approved small-molecule drugs commonly used in the clinical treatment of breast cancer, which were categorized into three primary classes and 20 subclasses, as detailed in **Table S1B**, and involved in various signaling pathways (**Figure S1A**). Based on a previous drug combination screening study ^16^, we selected 914 combination-cell line tuples across 18 common breast cancer cell lines. Each drug pair included one drug at two different concentrations to achieve 50-90% cell viability for each cell line, alongside a second drug at a single concentration ^16^. Additionally, we incorporated 98 anti-cancer compounds (**Table S1B**), which were either FDA-approved, in clinical trials, or under investigation for breast cancer, pancreatic cancer, colon cancer, and lung cancer ^16^.

**Figure 1.**
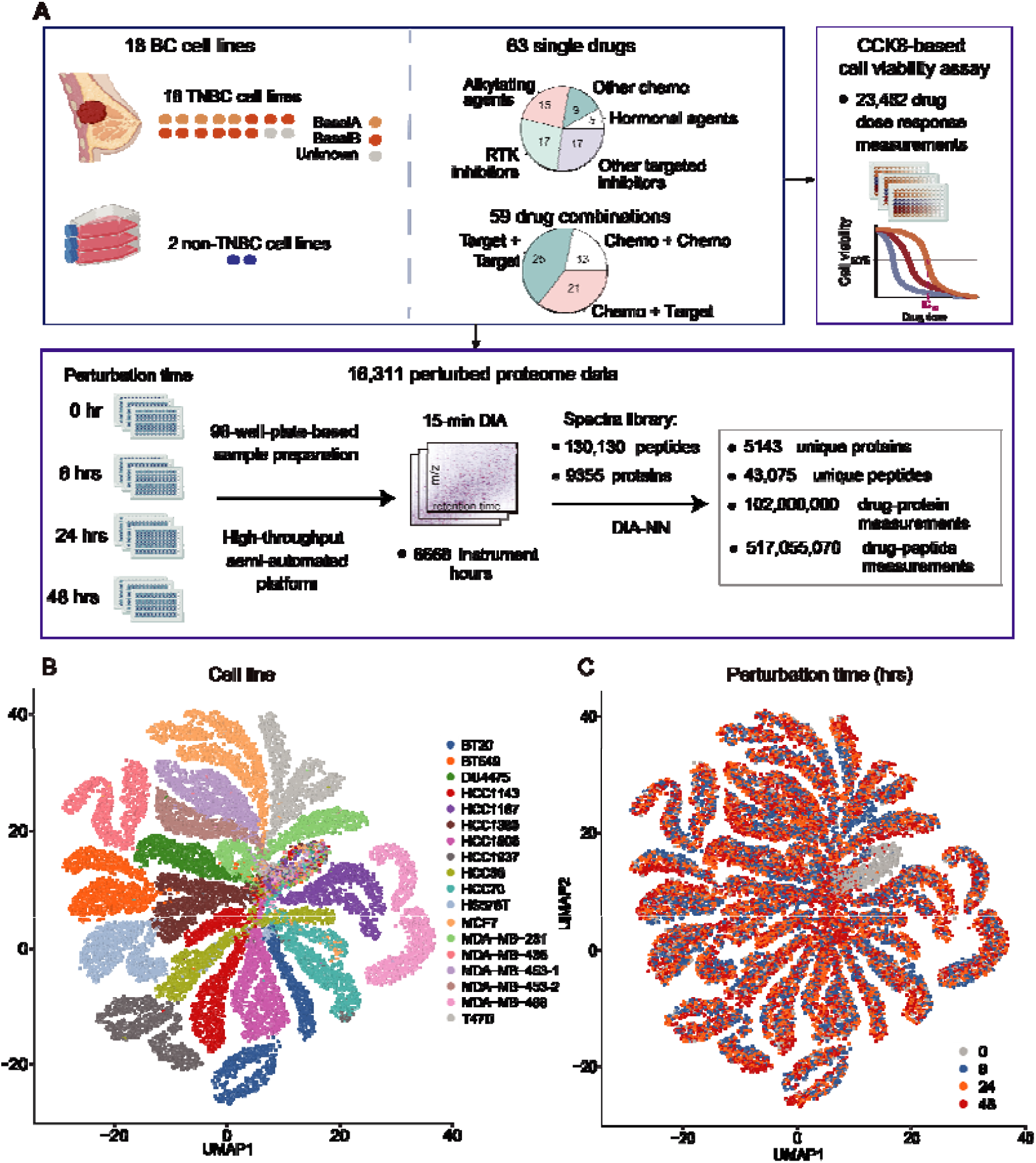
Overview of this perturbation proteomics study. **(A)** Study Design: Our perturbation proteomics approach involved an extensive panel of 18 cell lines, including 16 triple-negative breast cancer (TNBC) cell lines and 2 non-TNBC cell lines, which were exposed to a battery of 63 FDA-approved drugs over various time spans. Each condition was biologically replicated thrice, yielding a dataset encompassing 16,311 proteomic profiles and 23,554 cell viability assessments. **(B)**-**(C)** UMAP shows the unsupervised clustering of the whole proteomes by different cell lines **(B)**, perturbation durations **(C)**.

We treated the 18 cell lines with 63 FDA-approved drugs at 6, 24, and 48 hours, each in triplicates (**Figure 1A**). Approximately 10% of randomly selected samples were injected in replicates for quality control. Multiple control samples were analyzed to assess and correct batch effects arising from sample preparation and DIA-MS analysis. Altogether, we acquired 16,311 perturbation proteomic DIA-MS data files with approximately 6668 hours of mass spectrometry instrument time, yielding over 38 million high-quality perturbed protein measurements (**Figure 1A**). With DIA-NN analyses, the data led to the relative quantification of 5530 protein groups, corresponding to 5143 unique proteins (**Table S2A**). Data quality analyses are presented in **Figure S1**. Our proteomics dataset includes 252 proteins with known specific relevance to TNBC biology when compared to non-TNBC cells (**Table S2B**), 100 proteins characteristic in the basal A subtype, and 14 proteins specific to the basal B subtype (**Table S2C**), based on the seminal study conducted by Neel and colleagues ^17^. In addition, we collected 23,482 data points from cytotoxicity assays to complement the proteomic analysis. The quality control analysis demonstrated consistent reproducibility, validating the inclusion of protein measurements from biological replicates (**Figures S1B-C**). Our large-scale proteomic analysis revealed distinct differences among cell lines (**Figure 1B**), while variations in drug treatment durations (**Figure 1C**) and drug interference (**Figures 1D, S1E**) showed minimal impact on proteomic profiles. The resulting proteome datasets, termed ProteinTalks datasets (PTDS), are available at db.prottalks.com.

### The perturbation proteomics datasets capture drug-induced modulation of protein networks

To discern whether perturbed proteomes might signify the MOA for drugs, we firstly focused on the proteins directly targeted by the compounds. We identified 61 protein targets, accounting for 48.8% of the targets of the 63 drugs under investigation (**Figure S2A**).

Intriguingly, in cells resistant to capecitabine, thymidylate synthetase (TYMS) – a known target of the drug – demonstrated a substantial increase throughout the treatment duration (**Figure S2B**). To verify its role in drug resistance, we knocked down TYMS in the HCC1143 cell line and observed enhanced sensitivity to capecitabine (**Figures S2B-C**). The data imply that our perturbation proteomics data can illuminate potential molecular mechanisms responsible for drug efficacy and resistance.

Beyond the proteins directly targeted by the drugs, we also detected broader proteomic changes, indicating the indirect effects of drug treatment. We developed a quantitative metric called PertScore that encapsulates the aggregate prevalence of protein changes elicited by various perturbation conditions, encompassing drug types, treatment durations, and cell lines (see **Methods**). Using PertScore, we could pinpoint the most recurrent protein alterations stemming from drug perturbations (**Figure S2D**). We identified a total of 1123 perturbation-related proteins (PertScore > 10) (**Figures S2E, Table S3**). Upon enriching pathways, we observed that MOA-associated pathways were significantly perturbed at 6 hours, while cell death-related pathways were enriched at 48 hours (**Figure 2A**). Our data suggest that proteins in certain pathways act as vanguards for phenotypic shifts in cancer cells. For example, proteins differentially expressed due to alkylation agents are predominantly enriched in the DNA damage repair pathway, while hormone-based treatments result in changes primarily in the lipid metabolism pathway. Proteins affected by microtubule inhibitors show enrichment in the cellular cytoskeleton and associated pathways. Moreover, proteins with altered expression resulting from CDK inhibitor treatment are enriched in cell cycle-related pathways.

**Figure 2.**
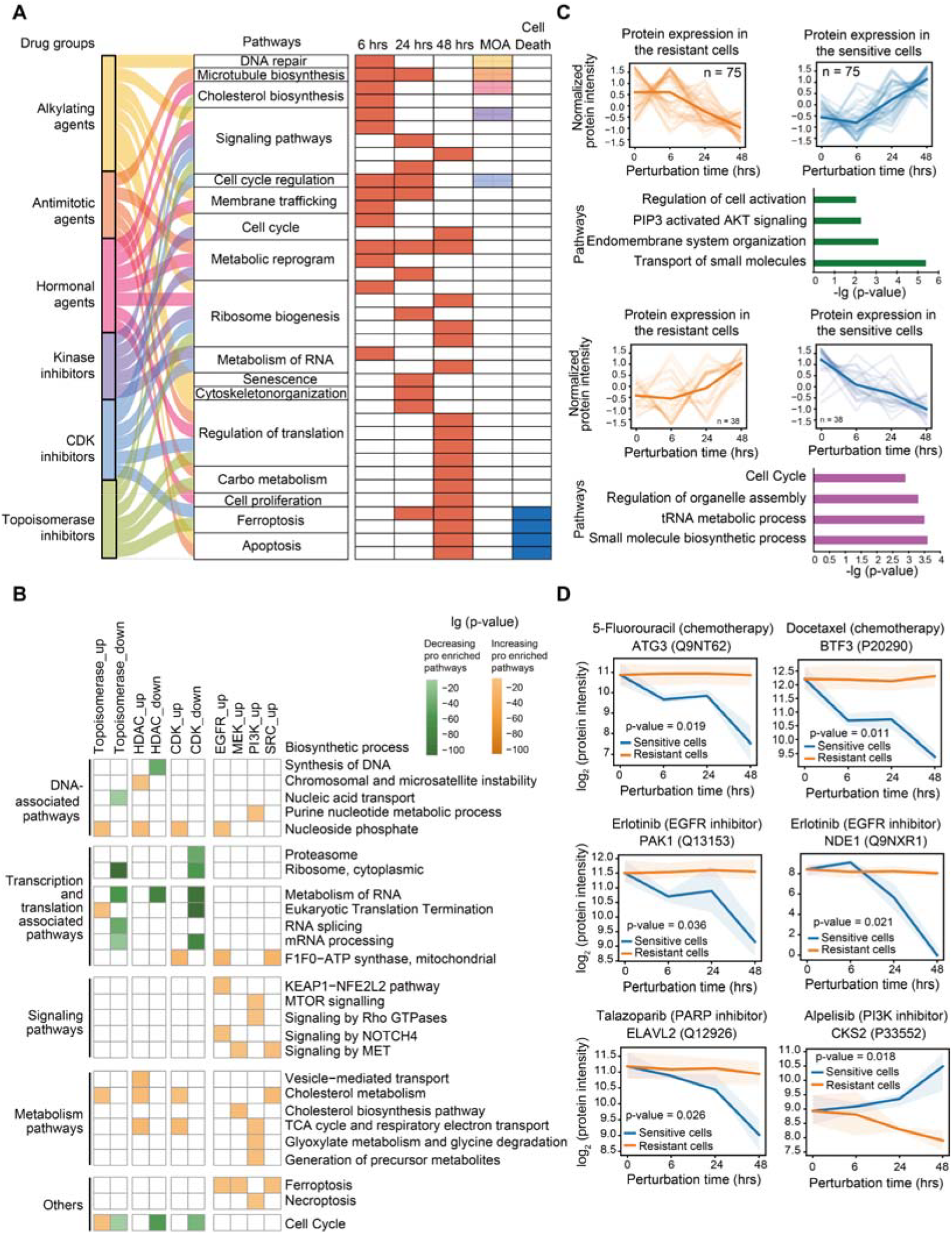
Longitudinal proteomic responses to drug perturbation. **(A)** The top five pathways enriched by differentially regulated proteins, as determined by PertScore (see **Figure S2D**), over various treatment intervals. Pathways associated with the mechanism of action (MOA) and cell death were specifically annotated. **(B)** Pathways enriched by proteins that were consistently upregulated or downregulated post-drug perturbation. Pathways associated with proteins exhibiting a decrease in abundance are represented in green, while those associated with proteins with increased abundance are shown in orange. **(C)** Clusters of proteins exhibiting sustained dysregulation throughout the duration of chemotherapy drug exposure, with an emphasis on reversed trends between resistant and sensitive cell groups. The enriched pathways related to these protein clusters are also depicted. **(D)** Temporal changes in protein levels that were distinctively expressed between resistant and sensitive cell groups. The p-value indicates the significance of the interaction between time points and the response of sensitive versus resistant cells, as assessed by two-way ANOVA.

Next, we narrowed down to the consistently dysregulated proteins resulting from the drug subtype perturbations through mFuzz analysis. Our findings indicate that both chemotherapy and targeted drugs increase the expression levels of proteins related to nucleic acid synthesis and fatty acid metabolism, while decreasing the expression levels of proteins involved in RNA processing, RNA metabolism, and cell cycle progression (**Figure 2B**). The ascending proteins along the treatment of kinase inhibitors are linked to various kinase-associated signaling pathways such as MET, NOTCH4, and mTOR. Diverse cell death pathways are also upregulated with drug treatment (**Figure 2B**). Therefore, each compound interacts with one or more protein targets and induces proteome-level changes via both common and perturbagen-specific modulation of cellular processes that reflect its MOA. These perturbagen-induced abundance changes vary in magnitude from protein to protein.

### Proteome dynamics is associated with drug resistance

Furthermore, we explored longitudinal changes related to drug sensitivity within this dataset. A total of 113 proteins exhibited differentially dynamic expression in response to drug treatment, effectively distinguishing between the chemotherapy-resistant and sensitive groups (**Table S3**). Specifically, 75 proteins exhibited an increase in expression following perturbation in the resistant group but decreased in the sensitive group (**Figure 2C**). These proteins were mainly enriched in cell cycle associated pathways, protein localization, purine ribonucleotide metabolic process, and carbon metabolism (**Figure 2C**). Conversely, 38 proteins showed an inverse dynamic expression pattern and were mainly associated with small molecule transport, endomembrane system organization, and the AKT signaling pathway (**Figure 2C**).

Multiple proteins maintained high expression levels in the resistant group while decreased in the sensitive group (**Figure 2D**), suggesting their involvement in drug resistance. For example, *ATG3*, an autophagy-related gene, showed a different dynamic trend between the two groups in response to 5-Fluorouracil, an anti-metabolic agent (**Figure 2D**). ATG3 has been implicated in chemotherapy resistance in HCC treatment by regulating autophagy processes ^18^. BTF3 also exhibited a distinct pattern in expression between two groups treated with Docetaxel, an anti-mitotic agent (**Figure 2D**). The upregulation of BTF3 indicates that the epithelial cancer cells may possess stemness characteristics ^19^, while cancer stem cells have shown chemotherapy resistance ^20^. PAK1 and NDE1 manifested this different dynamic trend after perturbation of Erlotinib, an EGFR inhibitor (**Figure 2D**). PAK1 has been associated with resistance to tyrosine kinase inhibitors in EGFR mutant lung cancer ^21^. NDE1 has been verified to interact with EGFR ^22^, indicating that the expression of NDE1 might influence the efficacy of the EGFR inhibitor Erlotinib (**Figure 2D**). ELAVL2 responded differently in the two groups when treated with Talazoparib, a PARP inhibitor (**Figure 2D**).

ELAVL2 has been implicated in drug resistance through the regulation of glycolysis ^23^. Conversely, the expression of CKS2 was elevated in the sensitive group while decreased in the resistant group (**Figure 2D**), highlighting its opposite effect on drug resistant compared with the previously mentioned proteins. Consistent with the published study, CKS2 exerts an antagonistic effect on the PI3K/Akt pathways ^24^.

### Development of a dynamical foundation model ProteinTalks

After benchmarking the proteomics dataset, we then established a dynamical foundation model, namely ProteinTalks, to achieve systematic understanding of protein network dynamics using an ordinary differential equation (ODE) integrated with the perturbation-aware neural network. We pretrained ProteinTalks using a dataset of 38 million protein measurements from 16,311 perturbed proteomic data and the characteristics of the drugs. The model contains two modules (**Figure 3A**). The first module incorporates baseline proteome data from untreated cell lines and drug targets. Through an encoder, the model is trained to predict perturbed proteomes at multiple time points. These predictions are then compared to the ground-truth proteome data, resulting in the calculation of a mean squared error (MSE) loss, referred to as Loss_1_ (**Figure S3**). This module stores the information of the protein network dynamics. In the second module, the predicted perturbed proteomes, combined with the structure of these drugs, including 881-dimensional drug molecular fingerprints (DMF), 55-dimensional drug physicochemical properties (DDP) and 61 targets of 63 drugs, are used to learn the cellular response to various perturbagens and the core proteins associated with drug response through a multilayer perception (MLP) (**Figure 3A, S3)**. More technical details of the ProteinTalks model construction are provided in **Methods**. This ProteinTalks foundation model enables a wide range of applications in drug discovery and precision medicine.

**Figure 3.**
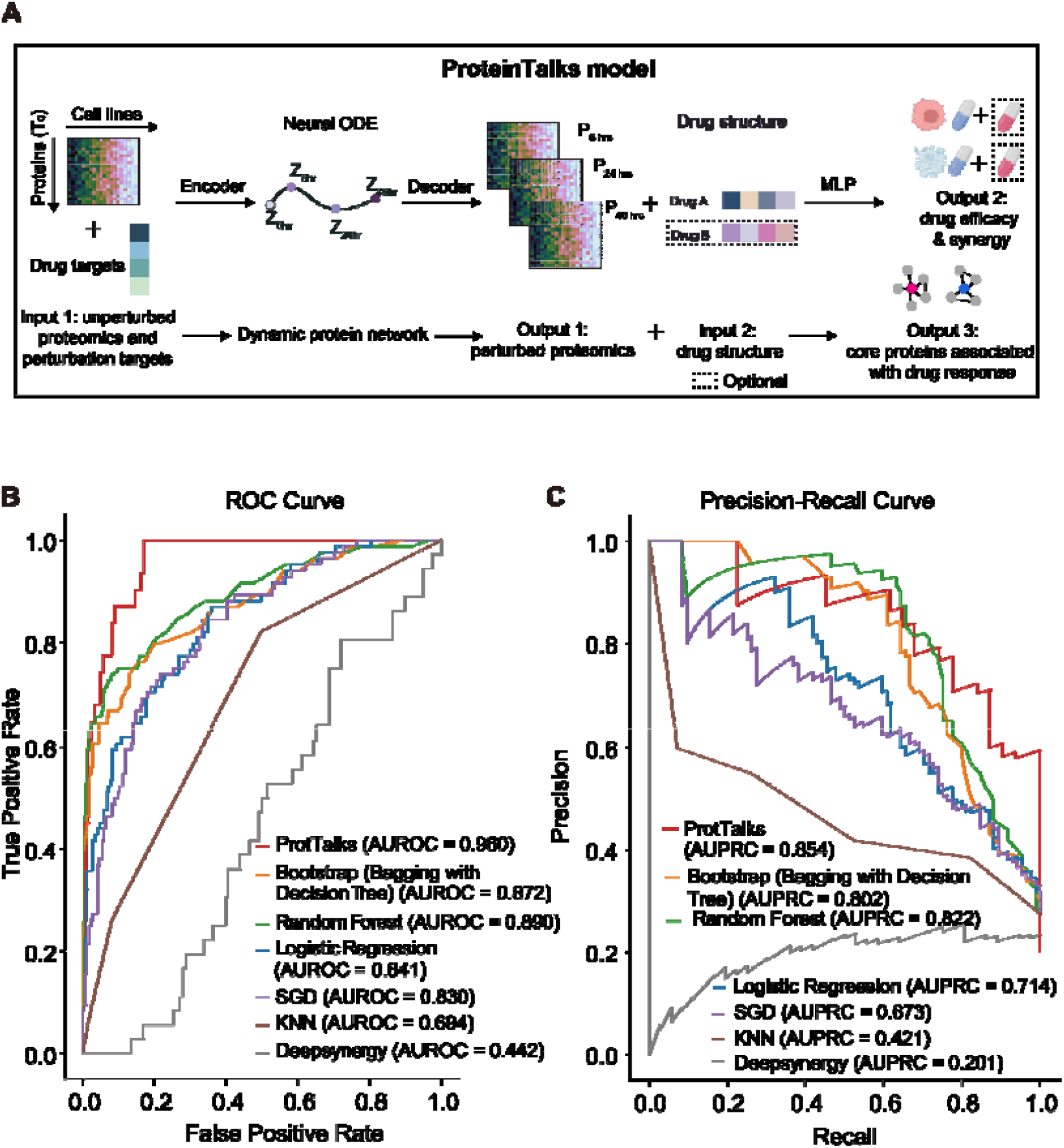
Development and performance of the ProteinTalks model. **(A)** The architecture of ProteinTalks. The baseline proteome from untreated cell lines and the list of drug targets are encoded into ProteinTalks, a linear network, to predict the perturbed proteomes, which are then decoded. Then the predicted proteome output from the first module, along with the initial time point proteome data and drug features obtained from SMILES descriptors, are processed through a linear layer to predict the essential proteins associated with drug response and effectiveness of single drugs or drug combinations. MLP, multilayer perception. **(B)-(C)** AUROC **(B)** and AUPRC **(C)** of six models with the ProteinTalks model.

### Prediction of responsiveness of drugs

We tested whether ProteinTalks model could boost the prediction of drug responsiveness through protein network dynamics in the biological systems based on unperturbed omics data. Notably, the ProteinTalks outperformed typical machine learning models, including bootstrap, random forest, logistic regression, SGD, KNN, and DeepSynergy (area under the receiver operating characteristic curve (AUROC) = 0.960, area under the precision-recall curve (AUPRC) = 0.854, accuracy = 0.910) (**Figure 3B-C**).

To avoid potential overfitting, we designed a leave-one-cell line-out cross-validation to test the ProteinTalks model (see model setting 2 in **Methods**). For the majority of cell lines (16 out of 18), the AUROC values were at least 0.9. Additionally, 16 out of 18 cell lines demonstrated the AUPRC values of at least 0.8 (**Figure S4**).

The chemical structure space is almost infinite; therefore, it is crucial to determine whether our model works in chemicals absent in the training data. We then performed leave-one-drug-out cross-validation for the ProteinTalks model (see model setting 3 in **Methods**). We stratified the drugs into four classes based on their accuracy, AUPRC, and AUROC values computed by ProteinTalks (**Figure S5A**).

The first class of drugs, which includes 14 drugs, showed relatively high accuracy and high AUPRC/AUROC values. The second class, comprising 21 drugs, exhibited relatively high accuracy but low AUPRC/AUROC values, due to an imbalance in the numbers of ineffective and effective drugs for each cell line. Specifically, the ratio of ineffective to effective among these 21 drugs was significantly higher compared to the other 42 drugs (**Figure S5B**). The third class, including 15 drugs, showed relatively low accuracy but high AUPRC/AUROC, suggesting that while the model performs well, setting the classification threshold of the predicted score at 0.5 may not be optimal. For instance, the accuracy of toremifene citrate efficacy prediction across different cell lines decreased as the threshold of predicted score increased from 0 to 0.25, and then remained at zero for thresholds above 0.25 (**Figure S5C**). The last class, consisting of 13 drugs, displayed the worst performance with low accuracy and low AUPRC/AUROC values, likely due to the absence of drugs with similar MOAs in the training set. For example, three drugs, namely sonidegib diphosphate, irinotecan hydrochloride, and pemetrexed disodium hydrate, modulate protein targets (SMO, TOP1, and DHFR, respectively) that are absent in the training set.

We further generated a dataset to independently assess the model (**Figure S3**, model setting 3 in **Methods**). Here, we extended the perturbation to 98 additional anti-cancer small-molecular compounds, with the baseline proteomic data without perturbation and corresponding drug efficacy data. These compounds were used to treat four TNBC cell lines included in the pretrained dataset. We acquired 5138 drug-dose response measurements (**Figure S3**). For the new compounds in the dataset, the overall accuracy was 0.619, with an AUROC of 0.671. After excluding the drugs with MOA categories absent in the training set, both accuracy and AUROC increased to 0.844 and 0.840, respectively. Remarkably, the accuracies for drug MOA categories present in pretrained datasets were consistent, as illustrated in **Figure S5D**. This consistency across the two test datasets suggests the model’s generalization ability in predicting drug efficacy. The data collectively show that ProteinTalks can well predict the drugs with similar MOAs to those in the training set, while for drugs with distinct MOAs absent in the training set, it could still achieve an overall accuracy between 0.6-0.7.

### Prediction of drug synergy

Next, we explored whether ProteinTalks could predict drug synergies. Our pretrained datasets covered 914 combination-cell-line tuples, which have also been investigated by the Garnett group at the Wellcome Sanger Institute ^16^. The ProteinTalks model, taking the structure information of each drug pair as input, is capable of predicting the synergy of drug combination, as evaluated by the Garnett group’s drug synergistic effect (**Figure 3A**). Our model showed that the synergistic scores of synergistic pairs were significantly higher than those of non-synergistic pairs (**Figure 4A**). Among the top 1000 tuples, the majority of potent synergies (58.7%) were observed in combinations of targeted drugs. The next significant proportion (26.2%) consisted of combinations of targeted drugs with chemotherapies.

**Figure 4.**
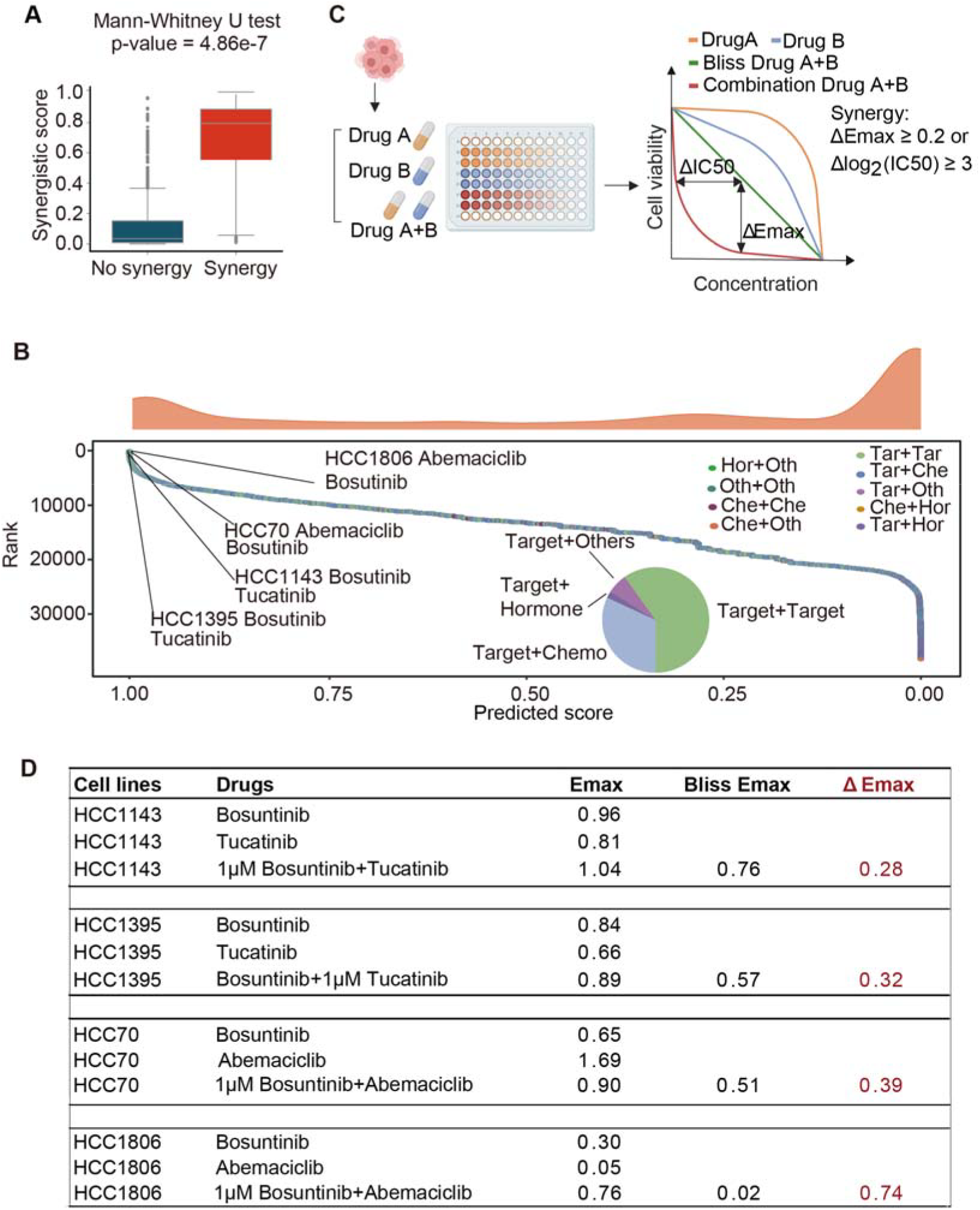
Deciphering the ProteinTalks Model’s Predictive Power for Drug Combination Synergy. **(A)** Deciphering the ProteinTalks’ Predictive Power for Drug Combination Synergy ^16^. Statistical significance was assessed using the Mann-Whitney U test. **(B)** Ranking of drug combinations based on their predicted synergy scores, with the highly synergistic combinations explicitly highlighted. The distribution of drug groups within the top 1000 synergistic combinations, as forecasted by the ProteinTalks model, is depicted in a pie chart. Tar, target drug; Che, chemotherapy drug; Hor, hormonal drug; Oth, other drug. **(C)** Description of the cytotoxicity assay protocol utilized for evaluating drug combinations and the specific criteria applied to determine drug synergy. **(D)** Results from cytotoxicity assays for drug combinations predicted by the ProteinTalks model to be synergistic, including combinations of bosutinib with tucatinib in HCC1395 and HCC1143 cells, and bosutinib with abemaciclib in HCC70 and HCC1806 cells. The treatments were administered as follows: HCC1143 cells received a combination of bosutinib and tucatinib with a fixed concentration of 1 μM bosutinib; HCC1395 cells were treated with the same drugs but with a fixed concentration of 1 μM tucatinib; and HCC70 and HCC1806 cells were exposed to a combination of abemaciclib and bosutinib with a fixed concentration of 1 μM bosutinib. Each assay plate included triplicate wells. The shifts in efficacy (ΔEmax), representing the reduced cell viability, were determined by calculating the difference in efficacy between the observed combination response and the expected response based on Bliss independence ^48^.

Combinations of targeted drugs with other types of drugs (12.3%) followed, while the least prevalent synergies (2.8%) were observed between targeted drugs and hormonal agents (**Figure 4B**). These findings suggest that combining targeted drugs with other agents tends to result in synergistic effects. Then we experimentally validated the synergistic efficacy of four synergistic tuples including bosutinib-tucatinib-HCC1143, bosutinib-abemaciclib-HCC70, bosutinib-tucatinib-HCC1395, and bosutinib-abemaciclib-HCC1806 using CCK-8-based cell viability assays. The assays employed a fixed concentration of one drug and a discontinuous 10,000-fold (seven points) dose-response curve of the other drug, following the methodology described previously ^16^. According to the synergistic screening criteria ^16^ (**Figure 4C**, Δemax ≥ 0.2 or Δlog_2_(IC50) ≥ 3), four of the top 20 predicted synergistic tuples demonstrated synergy, including the combination of bosutinib and tucatinib in HCC1395 (ΔEmax = 0.31), bosutinib and tucatinib in HCC1143 (ΔEmax = 0.28), bosutinib and abemaciclib in HCC70 (ΔEmax = 0.39), bosutinib and abemaciclib in HCC1806 (ΔEmax = 0.74), and were confirmed to be synergistic (**Figure 4C-D**). These results support the capability of ProteinTalks in drug synergy prediction.

### ProteinTalks prioritizes pathways and proteins networks responsible for drug resistance

To investigate whether ProteinTalks could identify the pathways or proteins associated with drug efficacy, we calculated the SHapley Additive exPlanations (SHAP) values to determine the importance of the proteins in the ProteinTalks predictions through comparing effective and ineffective groups based on individual drugs or drug combinations. A higher SHAP value suggests a stronger positive correlation between protein expression level and drug sensitivity. Firstly, we computed the average SHAP values associated with 41 hallmark pathways involving the proteins incorporated in the ProteinTalks models (**Table S4A**). Drugs with similar MOAs exhibited similar SHAP values in these pathways (**Figure 5A**). Pathways with high SHAP values for drug or drug class could be a potential indicator of drug responsiveness. For example, DNA repair pathways are highlighted as vital for predicting the sensitivity to alkylating agents, while the PI3K-AKT-mTOR signaling pathways are central to interpreting responses to PI3K and AKT inhibitors (**Figure 5A**). Similarly, the estrogen response pathway takes precedence in the context of aromatase inhibitors, which block estrogen or androgen production, underscoring its role in the drug MOAs (**Figure 5A**).

**Figure 5.**
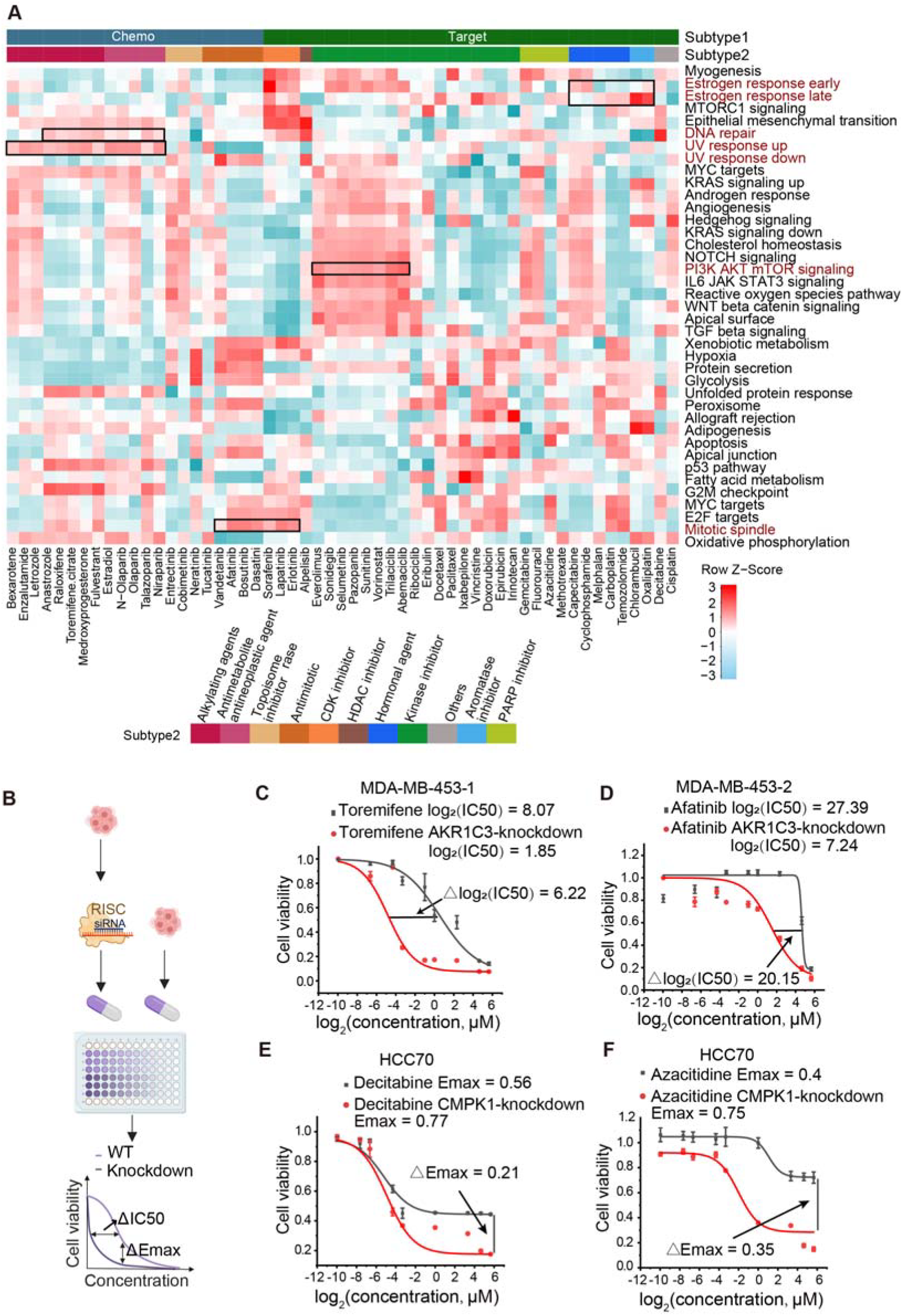
Exploring the Interpretability of the ProteinTalks Model. **(A)** Heatmap shows the SHAP values of each pathway for each drug after z-score normalization. Each row represents different protein pathways, and each column represents different drugs. Hierarchical clustering was performed using Euclidean distance and the complete linkage agglomeration method. Different colors in the columns indicate different categories of drugs. **(B)** Schematic of the cytotoxicity assay procedure, which incorporates siRNA-mediated gene knockdown in cell lines. **(C)** Outcomes of cytotoxicity assays on two MDA-MB-453 cell lines and HCC70 cells, post-transfection with AKR1C3 and CMPK1 siRNAs, followed by treatment with toremifene, afatinib, azacytidine, and decitabine, respectively.

We further extended the calculation of SHAP values to drug combinations, thereby enhancing the model’s interpretability. Within these combinations, the DNA damage and repair pathways played a critical role in accurately predicting synergy for chemotherapeutic combinations with other agents. Meanwhile, several signaling pathways, including PI3K-AKT-mTOR, TGF-β, and NOTCH, showed a high correlation with the responses to targeted therapies combined with other agents (**Figure S6**).

To evaluate the significance of these prioritized proteins in predicting drug sensitivity, we ranked the average SHAP value for each protein across different drug types. The top 30 proteins with the highest average SHAP values and the bottom 30 proteins with the lowest average SHAP values across all cell lines and drugs are displayed in **Figure S7**, providing a focused snapshot of the most important features. Among these, we observed numerous proteins typically targeted in breast cancer therapies, such as CDK4, CDK6, ERBB2, SRC, mTOR, and TOP2A, thereby validating the biological relevance of our model (**Figure S7**).

In addition to these established relevant proteins, our analysis also brought to light new ones. For instance, aldo-keto reductase 1C3 (AKR1C3) emerged as the most important protein for hormonal agents and the second-most important for kinase inhibitors (**Figures S7A-B**). Remarkably, AKR1C3 had the highest SHAP value in MDA-MB-453 cells treated with toremifene, an ER modulator, and also ranked first in terms of SHAP value in the context of treatment with afatinib, an EGFR inhibitor (**Table S4B**). AKR1C3 contributes to cell proliferation and differentiation by significantly enhancing the estrogen biosynthetic pathway^25^. Past studies have indicated that AKR1C3 overexpression may reduce TNBC cell sensitivity to doxorubicin ^26^. Among all samples treated with hormonal agents, the highest SHAP value of AKR1C3 ranking is MDA-MB-453 treated with toremifene (ER modulator) (**Table S4**). We further knocked down the expression of AKR1C3 using siRNA in the MDA-MB-453 cell line from two sources (**Figure 5D**). The efficacy of the siRNA knockdown experiments was confirmed with mass spectrometry analysis (**Figures S7D-E**). Cytotoxicity assays revealed that AKR1C3 knockdown enhanced the sensitivity of MDA-MB-453-1 cells to toremifene (Δlog2(IC50) = 4.76) and MDA-MB-453-2 cells to afatinib (Δlog2(IC50) = 3.32) (**Figure 5C**).

Within the alkylating agents category, TYMS was identified as the third most significant protein (**Figure S7C**), with its role in drug sensitivity corroborated in **Figures S2B-C**. Another protein involved in nucleic acid biosynthesis, CMPK1, ranked seventh. Knockdown of CMPK1 in HCC70 cells (**Figure S7F**) led to enhanced sensitivity to the kinase inhibitors decitabine (ΔEmax = 0.24) and azacytidine (ΔEmax = 0.26) (**Figure 5C**). In summary, the ProteinTalks model effectively uncovers interpretable proteins related to drug resistance.

### Finetuning allows drug response prediction based on transcriptome profiles of patient-derived xenografts in mice

Next, we explored whether the drug sensitivity prediction could be extended to breast cancer patient-derived tumor cells (PDTCs). We referred to a recent study of 30 short-term cultured PDTCs from patient-derived tumor xenograft (PDTX) models, treated with 96 compounds ^27^. Baseline transcriptome profiles of these PDTCs were acquired in this study. In the first-stage training, the ProteinTalks model was trained on varying ratios of the perturbation proteomic data (PTDS1-3 datasets), ranging from 0% to 90%. To address the differences between PTDS and PDTC datasets, we then implemented finetuning to build a multi-omics model, model-1. This process involved utilizing the 25 randomly selected baseline transcriptome profiles of PDTCs, phenotypic data of the 25 PDTCs’ response to 96 drugs, as well as drug information of 881 DMF, 55 DPP, and 40 targets (**Figure S8A**). As the PDTC dataset only provided drug efficacy data and lacked perturbed transcriptomic data, training Loss_1_ for model-1 was not feasible. The finetuning was conducted using Loss_2_ only (**Figure S8A**). For comparison, we trained the model-1 from scratch solely on the same subset of transcriptomic PDTC data without finetuning from the perturbation proteomic data, named model-1-wo (median accuracy: 0.786, median AUROC: 0.888, median AUPRC: 0.926). When 90% of the pretrained datasets were used in the first stage, the corresponding models obtained were model-1-90% (**Figure S8A**). The model-1-90% exhibited a significant enhancement in predicting of drug efficacy (median accuracy: 0.819, median AUROC: 0.918, median AUPRC: 0.951) (**Figure 6A, S8B**).

**Figure 6.**
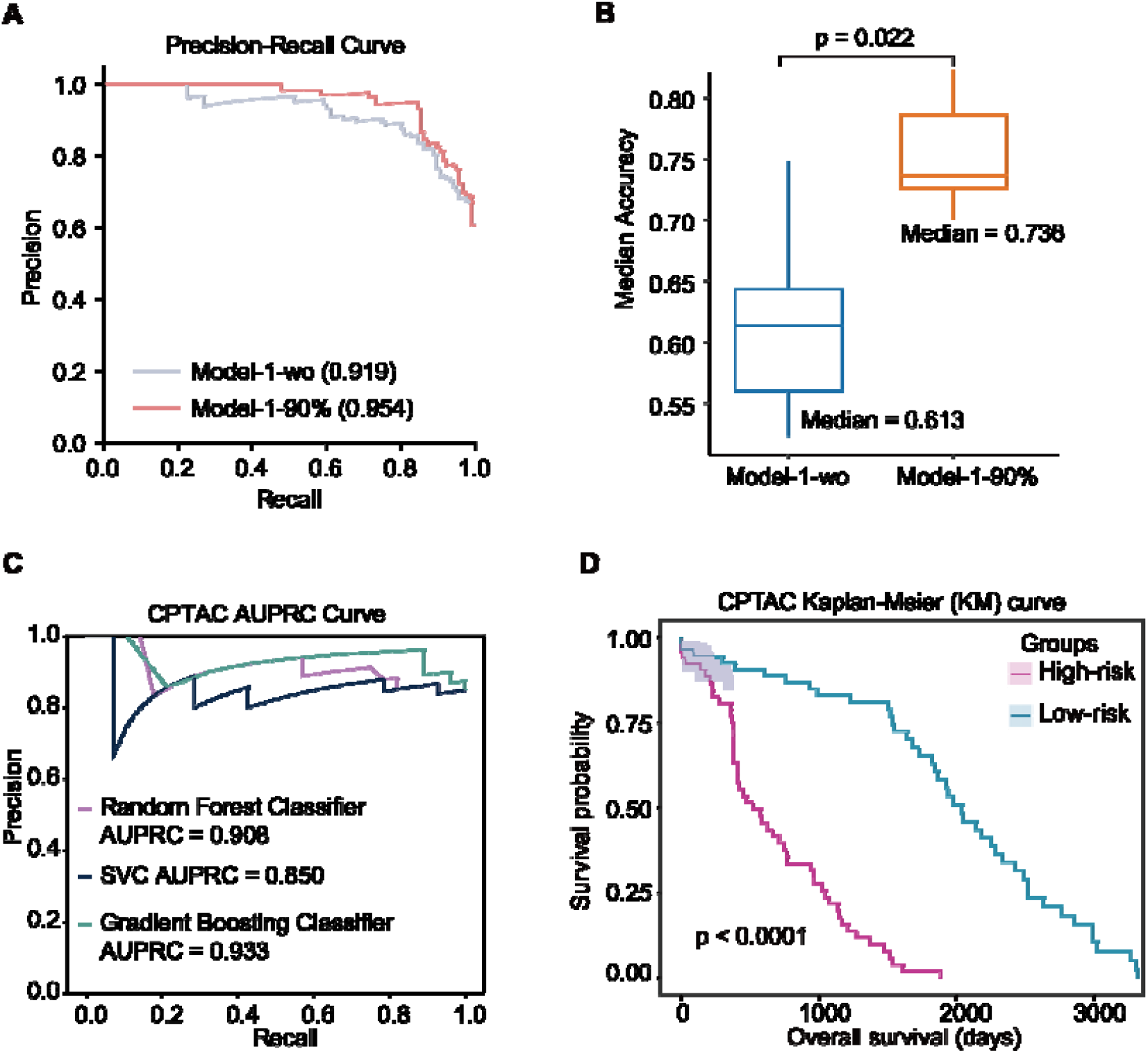
Assessing the Clinical Relevance of the Model-1. **(A)** In the BC patient-derived tumor cells (PDTCs)’ transcriptomic dataset, the AUPRC of the ProteinTalks model’s drug efficacy prediction performance between model-1-wo and model-1-90%. model-1-wo, model-1-without, was trained from scratch solely on the same subset of transcriptomic PDTC data without transfer learning from the perturbation proteomic data (**Figure S8A**). When 90% of the PTDS perturbation proteomic data was used in the first stage, the corresponding models obtained was model-1-90% (**Figure S8A**). **(B)** In the pan-cancer PDTXs’ transcriptomic dataset, the accuracy of the model’s drug efficacy prediction performance between model-1-wo and model-1-90%. model-1-wo, model-1-without, was trained from scratch solely on the same subset of transcriptomic PDTC data without transfer learning from the perturbation proteomic data (**Figure S8A**). When 90% of the PTDS perturbation proteomic data was used in the first stage, the corresponding models obtained were model-1-90% (**Figure S8A**). Statistical significance was determined via paired Welch’s t-test. **(C)** In the CPTAC proteomic dataset (N=107), the AUPRCs for the prognosis prediction performance in the test dataset (N=33) were evaluated using different machine learning models based on top the 60 proteins identified by ProteinTalks. **(D)** In the CPTAC proteomic dataset, the Kaplan-Meier (KM) curves of overall survival based on the top 60 proteins identified by ProteinTalks.

We also implemented similar finetuning on another 1075 pan-cancer PDTX models’ transcriptomic dataset to predict drug response ^28^. The median accuracy improved from 0.613 to 0.736 when comparing model-1-wo to model-1-90% (**Figure 6B**). This suggests the generalizability and adaptability of ProteinTalks in enhancing transcriptomic data for predicting therapeutic outcomes based on protein network dynamics.

### Application of ProteinTalks to clinical biopsy specimens

To further validate this approach using clinical biopsy tissue samples, we tested the drug response dataset in different terms. We firstly evaluated short-term drug response using baseline transcriptomic data from 16 clinical TNBC patients prior to receiving neoadjuvant carboplatin and docetaxel combination chemotherapy ^29^. Since our model can select core proteins associated with cellular response to specific categories of drugs, we assessed the effectiveness of the top 60 proteins screened by ProteinTalks in predicting chemotherapy sensitivity in these biopsy tissue samples. Specifically, the prognostic performance of these top 60 proteins was compared to that of the entire proteome and a randomly selected subset of 60 proteins. The results showed that the selected proteins significantly outperformed the random subsets in predicting chemotherapy drug sensitivity in TNBC and showed comparable or superior performance to the full protein set across multiple methods, including bootstrap, random forest, logistic regression, SGD, and KNN (**Figures S8C-E**).

We then performed long-term survival prognosis prediction in two public datasets: a transcriptomic dataset of 823 BC patients from TCGA (https://xenabrowser.net/) and a proteomic dataset of 122 BC patients from CPTAC ^30^. For these patients, who received various chemotherapy or targeted treatments in these datasets, we selected the top 60 proteins identified by ProteinTalks (**Table S5**), which represent the most affected by all categories of drugs. These proteins distinguished the patients into two groups, a high-risk group with poorer prognosis and a low-risk with better prognosis, in both TCGA (**Figure S9**) and CPTAC (**Figure 6C-D**) datasets.

These findings suggest that the ProteinTalks model effectively identifies clinically relevant biomarkers. It could improve the prediction of drug responses using baseline biopsies from TNBC patients prior to treatment, thereby potentially streamlining the pursuit of precision therapy.

## Discussion

Understanding the mechanisms and interactions within cells is a fundamental question in biology. To advance this, we generated a large-scale proteomic perturbation dataset and developed a corresponding dynamical foundation model named ProteinTalks, which also serves as a milestone towards constructing a virtual cell. The over 38 million protein measurements from 16,000 perturbed proteomic samples that we produced in-house makes it the largest proteomics dataset to date. In comparison, previous large-scale perturbation proteomics studies ^8,10,11,31^ were at least ten times smaller in size, insufficient to be used directly for building AI models.

Based on the large-scale temporal perturbation data, we developed a dynamical foundation model framework, which we believe is the first in its kind. core of the ProteinTalks model lies in describing the dynamic changes of variables over time, capturing the time-dependent behavior of the cell systems. This capability enables the model to gain a comprehensive understanding of protein network dynamics and can be applied in multiple downstream tasks. This model achieves an AUROC that is 5% to 27% higher than other models in predicting the efficacy of both single and combination therapies. Furthermore, we conducted validation experiments to confirm the synergy of four drug combinations. To delve deeper, we employed gene interference technology to confirm the involvement of specific proteins in drug response. For example, AKR1C3 was found to play a role in the response to toremifene and afatinib, while CMPK1 was found to contribute to the effects of decitabine and azacytidine. It creates a distinct latent space for clear differentiation between cellular states and drug types. Additionally, ProteinTalks can identify critical proteins and pathways associated with drug efficacy. For instance, DNA repair pathways play a significant role in predicting sensitivity to alkylating agents and chemotherapeutic combinations with other drugs. Similarly, the

PI3K-AKT-mTOR signaling pathways are essential for interpreting responses to PI3K and AKT inhibitors, as well as other targeted therapies when combined with other drugs. Through finetuning, this model enables predictions of the drug efficacy for patient-derived tumor xenograft (PDX) models. Furthermore, ProteinTalks can identify biomarkers associated with responses in 16 TNBC patients receiving neoadjuvant chemotherapy. Applied to multi-omics data of tumor samples collected before therapeutic treatment from TCGA and CPTAC, ProteinTalks can identify prognostic biomarkers in breast cancer patients, suggesting the potential of ProteinTalks to forecast treatment outcomes of diseases. Therefore, ProteinTalks represents a foundation model which has learned sophisticated protein network dynamics, and can be utilized to a broad range of downstream tasks.

Numerous AI models have been developed to map complex molecular networks for virtual cell construction. Among these, large-scale protein structure data have been used to build AlphaFold ^32,33^ for precise protein structure and interaction prediction. Single-cell RNA-seq data corpus have led to the development of multiple foundation models for cellular status prediction, such as Geneformer ^34^ and scGPT ^35^. However, there is no proteome foundation model for predicting protein network dynamics due to the limited availability of proteomic data, which constrains the application of AI in this context. Therefore, this is the first proteome foundation model, with dynamical information. Nevertheless, this study has several limitations. The proteome depth for each sample is less than several other recent papers ^8,10,11^. This is because we intentionally chose a high-throughput DIA-MS methodology of relatively low cost for this study (~70 RMB or 10 $ per proteome), otherwise, it won’t be feasible for a laboratory to acquire 16,000 perturbed proteomes to test whether this number of perturbations is sufficient for building a foundation model. Post-translational modifications (PTMs) are crucial for drug actions; however, we were not able to include them in this pilot study. Nevertheless, we have set up the framework of dynamical proteomic study in the context of foundation model and demonstrated its power in various tasks, paved way for future studies to put more efforts to identify more proteins, PTMs and drug combination perturbed proteomic data.

## Materials and Methods

### Cell line panel

BT20, BT549, DU4475, HCC1143, HCC1395, HCC1806, HCC1937, HCC38, HCC70, Hs578T, MDA-MB-436, MDA-MB-453-1, MDA-MB-468, HCC1187, T47D, and MCF7 cell lines were purchased from ATCC, while MDA-MB-453-2 and MDA-MB-231 were purchased from Meisen. The detailed information is listed in **Table S1A**. We have MDA-MB-453 cell lines from two sources. To distinguish them, the one from ATCC is denoted as MDA-MB-453-1, while the one from Meisen as MDA-MB-453-2.

### Cell viability assay

To determine the half maximal inhibitory concentration (IC50) in primary screening, we employed the CCK8 (Sigma) cell proliferation assay in accordance with the manufacturer’s protocol. Log-phase cells (**Table S1A**) were inoculated into a 96-well plate at a volume of 100 μL per well, using basal medium supplemented with 10% fetal bovine serum (FBS). After a 24-hour incubation, the medium was replaced with a gradient of drug concentrations (0.5, 5, 50, 200 µM) in medium containing 5% FBS, followed by another 24-hour incubation period (**Table S1C**). Cells without drug treatment served as negative controls. DMSO levels were ensured not to exceed 0.5%. Wells lacking cells acted as blank controls. Subsequently, 10 μL of CCK8 reagent (5 mg/mL) was added to each well, and the plates were incubated for 4 hours at 37°C. Absorbance was measured at 450 nm using a Multiscan Spectrum (BioTek, USA) to calculate the IC50 values. The criteria for drug efficacy were as follows:

1. Cell viability after drug treatment should increase with decreased drug concentrations (variance <20%) to be considered valid data. If cell viability rates treated with 0.5, 5, and 50 µM drug concentrations all exceed 50%, the drug is deemed ineffective.
2. If cell viability at 50 µM is not only less than 50% but also shows a further reduction to below 50% of cell viability observed at 0.5 µM, the drug is deemed effective, and vice versa.

Based on these criteria, we identified effective drugs and subsequently tested them across nine gradient concentrations over a 72-hour duration.

### Drug treatment for proteomics preparation

For cell culture and drug treatment, each 96-well plate included a control sample (no drug treatment) and a blank sample (no cells, medium only) for comparison. Samples within a single 96-well plate were treated as a batch. Triplicate biological replicates were performed for each treatment condition. The efficacy of each drug (IC50) was assessed bi-monthly on a minimum of two cell lines to ensure reproducibility. In single-drug experiments, a standard concentration of 10 µM was selected for perturbation, drawing on precedents from large-scale drug screening studies ^8,36,37^. A comprehensive assay of 2025 drug combinations ^16^ informed the concentrations used in combination treatments. Following drug treatment, cells were washed three times with PBS and lysed in lysis buffer containing 6 M urea, 2 M thiourea, and 10 mM ammonium bicarbonate buffer (ABB). Proteins were then reduced and alkylated using TCEP and IAA, respectively, followed by digestion with trypsin.

### LC-MS/MS analysis

For the proteomics analysis, we injected 1 µg of purified peptides into an Eksigent Nano LC 415 system (with a 1-10 µL/min flow module to switch the LC from nano-flow to micro-flow). Chromatographic separation was achieved using an Eksigent analytical column (C18 ChromXP, 3 µm, 0.3 x 150 mm) and trap column (C18 ChromXP, 5 µm, 0.3 x 10 mm), as detailed in prior documentation ^38^. Mobile phase Buffer A consisted of 2% acetonitrile (ACN) with 0.1% formic acid (FA), and Buffer B comprised 98% ACN with 0.1% FA. A linear gradient of 5-32% Buffer B was applied over 15 minutes at a flow rate of 5 µL/min. The coupled DIA-MS analysis was conducted on a TripleTOF 5600+ system, with the ion accumulation time set at 150 ms for MS1 (m/z 350-1250) and 20 ms for each DIA window. We optimized the DIA window scheme to include 71 variable windows and operated the instrument in high sensitivity mode.

To generate the spectral library, we fractionated 10 mg of peptides derived from various TNBC cell lines using the Ultimate 3000 system, following established protocols ^39^. The resultant TNBC-specific spectral library encompassed 130,130 peptides corresponding to 9355 proteins.

In DIA-MS analysis, a pooled sample of nine TNBC cell lines (BT20, BT549, HCC1143, HCC1395, HCC1806, HCC1937, HCC38, HCC70, Hs578T), along with randomly chosen technical replicates, were employed as quality control samples within each batch.

Raw DIA data files were initially converted to profile mode mzML format using msConvert. Subsequent analysis of mzML files was performed with DIA-NN software (version 1.7.15), as previously described ^40,41^. Analysis parameters were meticulously set: trypsin/P as the protease with a maximum of two missed cleavages allowed; N-terminal methionine excision, cysteine carbamidomethylation, and methionine oxidation as the specified modifications; peptide length restricted to a range of 7 to 30 amino acids; and an m/z range of 400 to 2000 for precursor ions and 100 to 2000 for fragment ions. False discovery rate (FDR) cutoffs for precursor ions, peptides, and proteins were stringently maintained at 1%. Data were processed using the single-pass mode of the neural network classifier.

### Proteomics data preprocessing and normalization

From the initial dataset, 689 samples with identification counts below 1000 were discarded, leaving 16,311 samples for analysis. No proteins exhibited a missing rate greater than 90% in the complete dataset, with an overall missing rate of 51.7%. The initial step in data preprocessing involved imputing missing values with a value equivalent to 0.8 times the minimum detected intensity. The reproducibility of biological replicates, technical replicates, and pooled samples was assessed using Pearson correlation and the coefficient of variation (CV) as metrics.

### Differentially expressed analysis

Prior to differential expression analysis, proteins that were absent in more than 80% of the samples across both groups were excluded. The comparison between the two groups was conducted using a two-sided unpaired Welch’s t-test. P-values obtained from the statistical tests were adjusted for multiple comparisons using the Benjamini-Hochberg (B-H) method. Differentially expressed proteins were identified based on a combination of fold change and either raw p-values or B-H adjusted p-values.

### PertScore calculation

The PertScore quantifies the cumulative frequency of protein expression alterations induced by various drugs across different time points in each cell line, aiming to highlight the most recurrent protein expression changes triggered by drug perturbations. Proteins were scored based on their perturbation impact and visualized in a three-dimensional coordinate system (Figure S2A), where the x-axis represents different cell lines, the y-axis denotes drug types, and the z-axis indicates drug treatment durations. Each coordinate (Xl, Ym, Zn) signifies the outcome of the differential analysis between samples from cell line l treated with drug type m at time point n and their corresponding untreated controls. For a given coordinate, if a protein is significantly upregulated, it is assigned a score of 1. Conversely, a significant downregulation is scored as −1. Proteins that do not exhibit significant changes are assigned a score of 0.

### Pathway enrichment

To elucidate the biological pathways associated with the differentially expressed proteins, we employed four well-established databases: KEGG pathway ^42^, Metascape ^43^, STRING ^44^, and Ingenuity Pathway Analysis ^45^ (IPA, version 51963813). Pathway enrichment analysis was performed with a defined significance threshold of p < 0.01, and only pathways containing at least two proteins or metabolites from our dataset were considered for further analysis.

### mFuzz analysis

One-way analysis of variance (ANOVA) was used to determine differences between samples treated at different time points (p < 0.05). The average normalized protein quantities, z-score in each GS grade were used for fuzzy c-means clustering with the R (version 4.0.2) package Mfuzz (version 2.48.0). The number of clusters was set to four, and the fuzzifier coefficient, M, was set to 1.25. The clusters with consistently increasing or decreasing trends were selected. In **Figure 2B**, the clustering was based on the drug group, while in **Figure 2C**, the clustering was based on both the drug group and drug sensitivity. The proteins that overlapped between the increasing clusters in the sensitive group and the decreasing clusters in the resistant group were filtered out. This filtration process is depicted in the upper panel of **Figure 2C**, and the same procedure was conducted in reverse for the other group.

### Data preprocessing

After obtaining data from different cell lines, various drug perturbations, and multiple time points, we averaged the samples from the repeated experiments, for example, data for 6 hrs in HCC70 with drug #75. Prior to inputting the data into the model, we performed min-max normalization on each sample. Ultimately, this process yielded the expression levels of 5585 proteins under each perturbation condition, across different cell lines, at 0, 6, 24, and 48 hours.

### SMILES generation

Using the SMILES representations of the drug molecules, we employed the R-package ChemmineR ^46^. We set the parameters as follows, functions MW and MF parameter addH=FALSE, function bonds parameter type=“charge”, function groups parameter type=“countMA”, function rings parameters upper=6, type=“count”, arom=TRUE. We obtained the 881-dimensional drug molecular fingerprints (DMF) and the 55-dimensional drug physicochemical properties for each drug.

### Data partitioning for model performance evaluation

We evaluated the model’s performance under three distinct datasets with different partitioning settings:

#### Setting 1: Model Performance Test

All time points are treated as a combined sample. The dataset, including various cell lines and drug perturbations, is randomly divided into training, validation, and test sets with a split ratio of 0.7:0.2:0.1, resulting in 1070, 305, and 154 samples respectively.

#### Setting 2: Model Performance Test for Cross-Cell-Type Data

One cell line is left out at a time. The model is trained and validated on data from the remaining cell lines and tested on the excluded cell line, assessing cross-cell-type performance.

#### Setting 3: Model Performance Test with Individual Drugs

Tests the model on previously unseen drugs using the results from Setting 1. This assesses the model’s generalizability to new drug perturbations.

These settings ensure a comprehensive evaluation of the model’s robustness and generalizability in predicting protein responses to drug perturbations.

### Model architecture and parameters

The initial component of our model is devoted to predicting post-perturbation protein expression using pre-perturbation protein expression as input data. Initially, the model ingests data from 5,585 proteins *P*_0_, concatenating this with corresponding dimension perturbation data *D*. This is processed through a linear layer *L*_1_, which elevates the dimensionality from 2 dimensions to 32 dimensions:

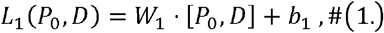

where *W*_1_ is the weight matrix and *b*1 is the bias vector.

Subsequently, a convolutional network C_1_ is employed to further enhance the dimensionality to 128 dimensions. Following this, we have engineered a two-layer linear network, utilizing the Softplus as the activation function and implementing a dropout ratio of 0.1. The output of the convolutional network is passed through two linear layers *L*_2_ and *L*_3_ :

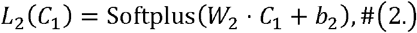

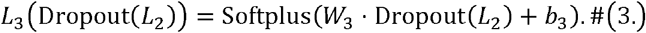

The next step involves parameterizing this two-layer linear network with Neural Ordinary Differential Equations (Neural ODEs), opting for the ‘rk4’ solver, a fourth-order Runge-Kutta method, to predict omics data over multiple time points:

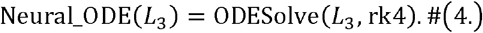

Then, a convolutional network *C*_2_ is applied to reduce the dimensionality of the predicted multi-time point embeddings, decoding them down to 32 dimensions. Finally, a linear layer *L*_4_ is utilized to decode the 32-dimensional representation back into the proteomics space, and get predicted proteomics 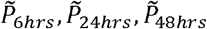:

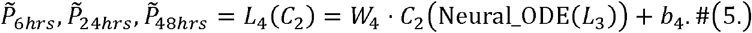

By utilizing an MSE loss, the first part of ProteinTalks denoted a loss function as Loss_1_:

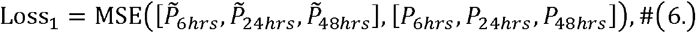

where P_6hrs_, *P*_24*hrs*_, *P*_48*hrs*_ is the ground-truth of proteomics at time 6h, 24h, 48, respectively.

The second part of our model is centered around the prediction of drug efficacy. This involves concatenating the proteomics data 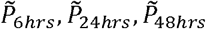 predicted at specific or all time points with the initial proteomics data *P*_0_, followed by a convolutional network *C*_3_ that elevates the data dimensionality to 128 dimensions:

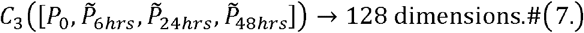

To describe the physicochemical properties of small molecule drugs, we utilize descriptions based on Simplified Molecular-Input Line-Entry System (SMILES), obtaining 935 features. These drug features for two drugs *D*_1_ and *D*_2_ (or duplicate if only one drug was used) are concatenated, forming a tensor [*D*_1_, *D*_2_] with a dimensionality of 935 x 2, which is then expanded to 128 dimensions through another convolutional network *C*_4_:

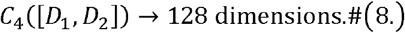

Subsequently, the proteomics data and drug features are concatenated. This combined dataset is processed through a linear layer *L*_5_ utilizing the ReLU activation function to reduce its dimensionality to 32 dimensions:

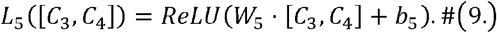

Finally, the model was followed by a linear layer, applying a sigmoid function to the output, which yields a prediction on the efficacy of the drug or drug combo synergy 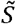:

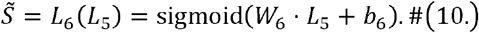

By utilizing a binary cross entropy (BCE) loss, the second part of ProteinTalks denoted a loss function as Loss_2_ :

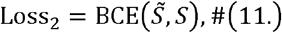

where *S* is the ground-truth of drug efficacy or combo synergy.

Final loss function for ProteinTalks is:

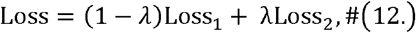

where *λ* is the hyperparameter for balancing the importance of the two tasks. The *λ* was set to 0.8 for ProteinTalks.

### Multi-task learning

Given that ProteinTalks is a multi-task learning task, we address the gradient conflicts between the losses from different tasks as Loss function (12), specifically Loss1 and Loss2, by calculating their cosine similarity. The cosine similarity of the gradients is computed as follows:

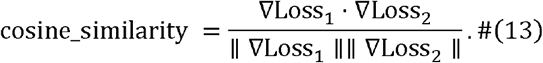

We then apply a clipping operation using a cutoff value of 1.0:

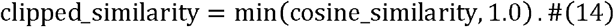

Based on the clipped cosine similarity, we adjust the weights of the losses by a factor of 0.01:

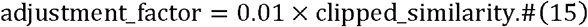

The total loss is updated by modifying the weight of each task based on the gradient similarities. If the gradient directions are similar or aligned, we retain the original weights. If the gradient directions conflict, we prioritize the drug efficacy prediction task by increasing its gradient weight while decreasing the weights of other tasks. The adjustment is done as follows:

If gradients are similar:

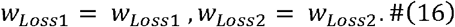

If gradients conflict:

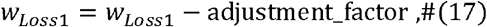

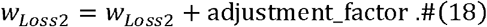

Finally, we ensure that the weights remain within a reasonable range. The updated gradient is then backpropagated through the model to update the parameters. For implementation details, refer to the code.

### Implementation of relevant methods for benchmarking

By concatenating data from multiple time points and combining it with drug data, we modified the input data dimensions of the DeepSynergy code to fit our dataset. DeepSynergy is a fully connected network that integrates omics data and drug information into the model to predict the efficacy of drug combinations. Other parameters were kept as original. We then applied DeepSynergy to our data. For Bootstrap, we used Bagging with a Decision Tree, implemented with *scikit-learn*’s *BaggingClassifier* function, setting parameters as *n_estimators=100 and bootstrap=True*. For Random Forest, we used *scikit-learn*’s *RandomForestClassifier* function, with default parameters from version 1.3.0 of *scikit-learn*. For Logistic Regression, we used *scikit-learn*’s *LogisticRegression* function, setting the parameter *max_iter=1000*. For stochastic gradient descent, we used *scikit-learn*’s *SGDClassifier* function, with the parameter *max_iter=1000*. For the K-nearest neighbor algorithm, we used *scikit-learn*’s *KNeighborsClassifier* function, with default parameters from version 1.3.0 of *scikit-learn*.

### SHAP value calculation

The significance of individual proteins in relation to sensitivity of single drug or the synergy of drug combination was determined using SHAP (SHapley Additive exPlanations) values for Loss_2_ ^47^. The analysis commenced by calculating the SHAP value for each protein, thereby assessing its specific contribution to the predictive model’s output. For each cell line and drug condition, the SHAP values of each protein were computed for sensitive drug or effective drug combinations, using the resistant drugs or no synergy combinations from other cell lines or drugs as background samples. This evaluation framework permitted the assessment of particular proteins’ roles in predicting the effectiveness of drugs or their combinations.

### SHAP value filtration

Proteins were then ranked according to their average SHAP values across all samples. The top 30 proteins with the most significant positive SHAP values were identified as having a favorable influence on sensitivity of single drug or the synergy of the drug combination, while the 30 proteins with the highest negative SHAP values, in absolute terms, were recognized as having an adverse effect. This selection of 60 proteins represents the most influential factors, thus forming the core set of features with the most substantial impact on the efficacy of single drugs or drug combinations.

By leveraging SHAP values in this manner, we pinpointed key proteins that modulate the response to single drugs or drug combinations, enhancing our understanding of their mechanistic roles and informing the refinement of predictive models for drug sensitivity or drug combination synergy.

### Validation of clinical data for top 60 proteins by SHAP values

After model training, we selected the top 60 proteins with the highest SHAP values for antimitotic drugs. For 16 individual clinical samples, we conducted 4-fold cross-validation and repeated the process 100 times for each method. For Bootstrap, we used *scikit-learn*’s *BaggingClassifier* function with the parameter *n_estimators=10*. For Random Forest, we used *scikit-learn*’s *RandomForestClassifier* function with the parameters *n_estimators=100 and criterion=‘log_loss’*. For Logistic Regression, we used *scikit-learn*’s *LogisticRegression* function with the parameter *max_iter=1000*. For stochastic gradient descent, we used *scikit-learn*’s *SGDClassifier* function with the parameters *max_iter=1000, tol=1e-3, and loss=‘log_loss’*. For the K-nearest neighbor algorithm, we used *scikit-learn*’s *KNeighborsClassifier* function with the parameter *n_neighbors=3*.

### Transfer learning for the transcriptomics data of PDTC model

To illustrate the complementary nature of proteomics data to transcriptomics data, we conducted a study comparing the effects of models pretrained on proteomics data versus those trained from scratch on transcriptomics data ^27,28^. We employed various proportions of proteomics data to train the ProteinTalks models in the first stage, specifically using 90% for training and 10% for validation. Subsequently, models trained on proteomics data were transferred to transcriptomics data, utilizing a dataset comprising 90% samples for training and 10% for testing. Since the ground truth only includes drug efficacy data and lacks omics data at future time points, the hyperparameter *λ* in the loss function was set to 1. Consequently, the loss function for ProteinTalks is defined as:

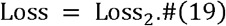

After training for 2000 epochs, the ProteinTalks model was obtained. The performance of these models was then assessed using the test dataset to validate the efficacy of leveraging proteomics data to enhance the predictive capabilities of transcriptomics-based models.

### Machine learning for the transcriptomics data

We extracted the expression matrix of these 62 proteins from the TCGA-BRCA dataset and CPTAC-BRCA dataset, and applied the train_test_split method of the scikit-learn package in Python to divide the dataset into the training dataset and the test dataset. Random forest, Support vector machine, and Gradient boosting decision tree models are used for training and prediction. The performances of three machine learning models were compared, and random forest showed the best performance on the test dataset.

For each model, we first built a GridSearchCV parameter search and then trained a classification model to predict alive/death. All models were trained using 3-folds cross validation in the train dataset (N =658) and tested against the test dataset (N =165). We found that random forest showed significantly better performance in independent validation sets. For the random forest model, our parameter adjustment range is: max_depth: [None, 2, 3], min_samples_leaf: [2, 4], min_samples_split: [2, 4, 8], n_estimators: [20, 30, 50, 100, 500, 1000]. The accuracy and AUPRC of the model on the test set are output.

### Survival analysis for the BC patients

Survival outcomes were defined using time-to-event data, with death as the endpoint and a binary incidentor for the event occurence. For the Cox model in the survival package, all 62 proteins were used as variables, and Cox multivariate analysis was performed to analyze the relationship between multiple proteins and sample survival in R. The Cox model calculated the coefficient value of each protein, which was then multiplied by the corresponding protein expression in each sample, and the average value was calculated as the risk score. The risk scores were divided into grouping variables using the surv_cutpoint function of survminer package. The Survfit function was then used to generate Kaplan-Meier survival curves.

### Gene interference in different cell lines

Breast cancer cell lines were subjected to gene interference using small interfering RNA (siRNA) in conjunction with Hieff Trans liposomal transfection reagent (Yeasen), following the manufacturer’s protocol. The siRNA sequences targeted against AKR1C3 and CMPK1 were as follows: for AKR1C3, 5’-GGAACUUUCACCAACAGAU-3’; and for CMPK1, 5’-GAGUAGUGGUAGGAGUGAU-3’. Initially, siRNA and transfection reagent were individually mixed with Opti-MEM medium (ATCC) and allowed to incubate for 5 minutes at room temperature. These mixtures were then combined and incubated for an additional 15 minutes at room temperature to allow complex formation. The siRNA-liposome complexes were added to the cell suspensions, which were subsequently transferred to 96-well plates for incubation over a 24-hour incubation period. Following transfection, the cells were processed for subsequent experimental steps, which included drug treatment for sensitivity assays and protein harvesting for proteomic analyses, as previously described.

## Acknowledgements

This work is supported by grants from Joint Funds of the National Natural Science Foundation of China (No. U24A20476), National Natural Science Foundation of China (Young Scientist Fund, No. 32401239), the Zhejiang Provincial Natural Science Foundation of China (LQ24C050002), Pioneer and Leading Goose R&D Program of Zhejiang (2024SSYS0035), National Key R&D Program of China (No. 2022YFF0608403, 2021YFA1301600), and National Natural Science Foundation of China (81972492). We thank Westlake University Supercomputer Center for assistance in data generation and storage, and the Mass Spectrometry & Metabolomics Core Facility at the Center for Biomedical Research Core Facilities of Westlake University for sample analysis. We thank Mr. Xuan Zhang, Ms. Mengmeng Liang, Ms. Xinxin Liu, Ms. Yinan Lai, Ms. Wangmin Lin, Ms. Yuan Hu, Ms. Mengni Chen, Mr. Chengliang Li, Dr Jianghai Wang, and Ms Yinfei Wu for assistance for the sample preparation and MS maintenance.

## Author contributions

T.G., H.W. and Y.Z., designed and supervised the project. R.S., L.Q., X.Z., W.H., Q.X., Z.L., and G.Z., conducted proteomic analysis. H.C, Z.X, L.T., and R.S. conducted bioinformatics analysis. Y.L., P.Z., and H.W. performed machine learning analysis. R.S., L.Q., Y.Z., and T.G. interpreted the data with inputs from all co-authors and wrote the manuscript with inputs from co-authors.

---

**Supplementary Figure 1.**
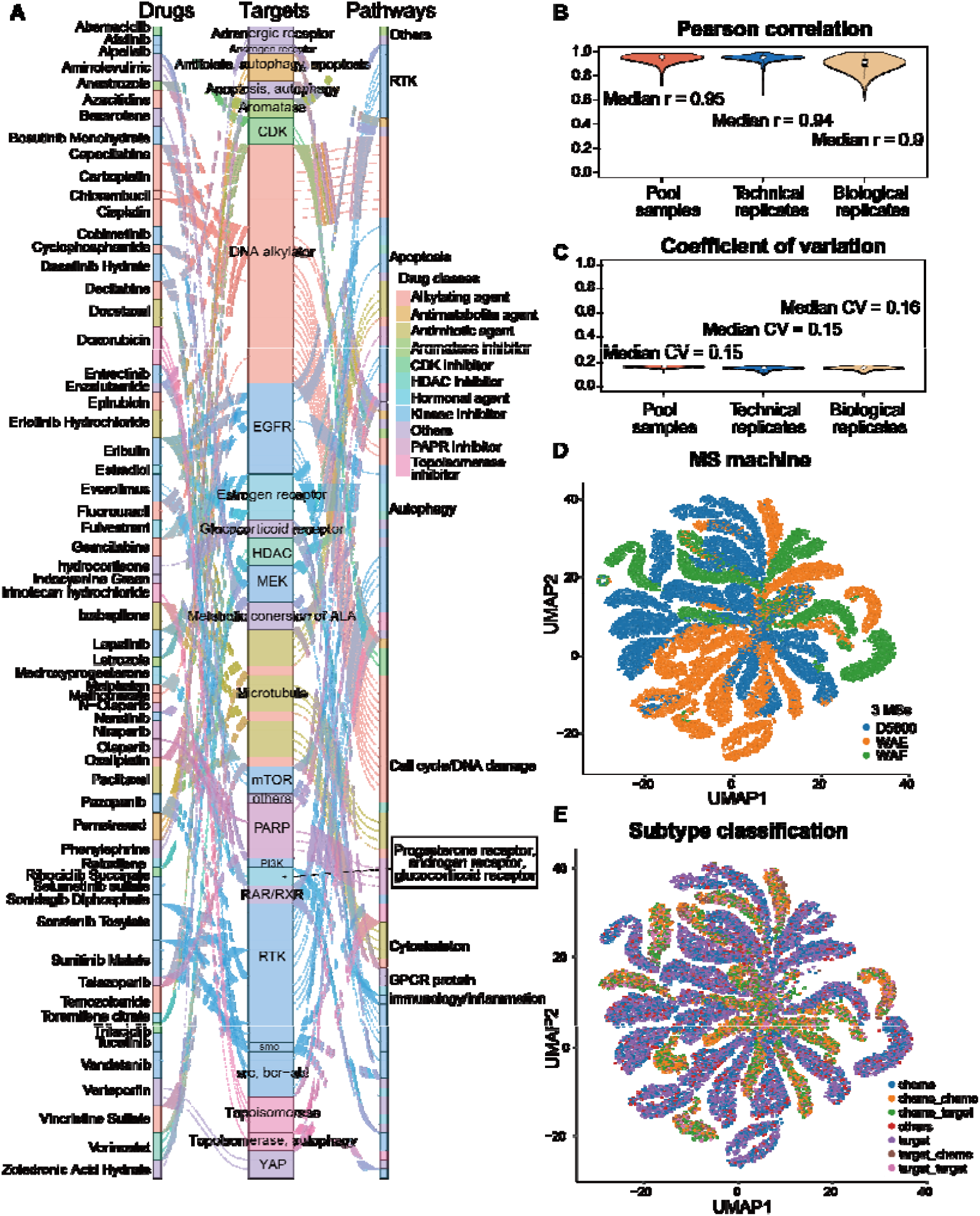
Quality control analysis. **(A)** Drug classification for 63 drugs. **(B)-(C)** The reproducibility of pool, technical replicate, and biological samples evaluated by Pearson correlation **(B)** and coefficient of variation **(C). (D)-(E)** UMAP shows whole proteomics grouping by MS machine **(D)** and drug classification **(E)**.

**Supplementary Figure 2.**
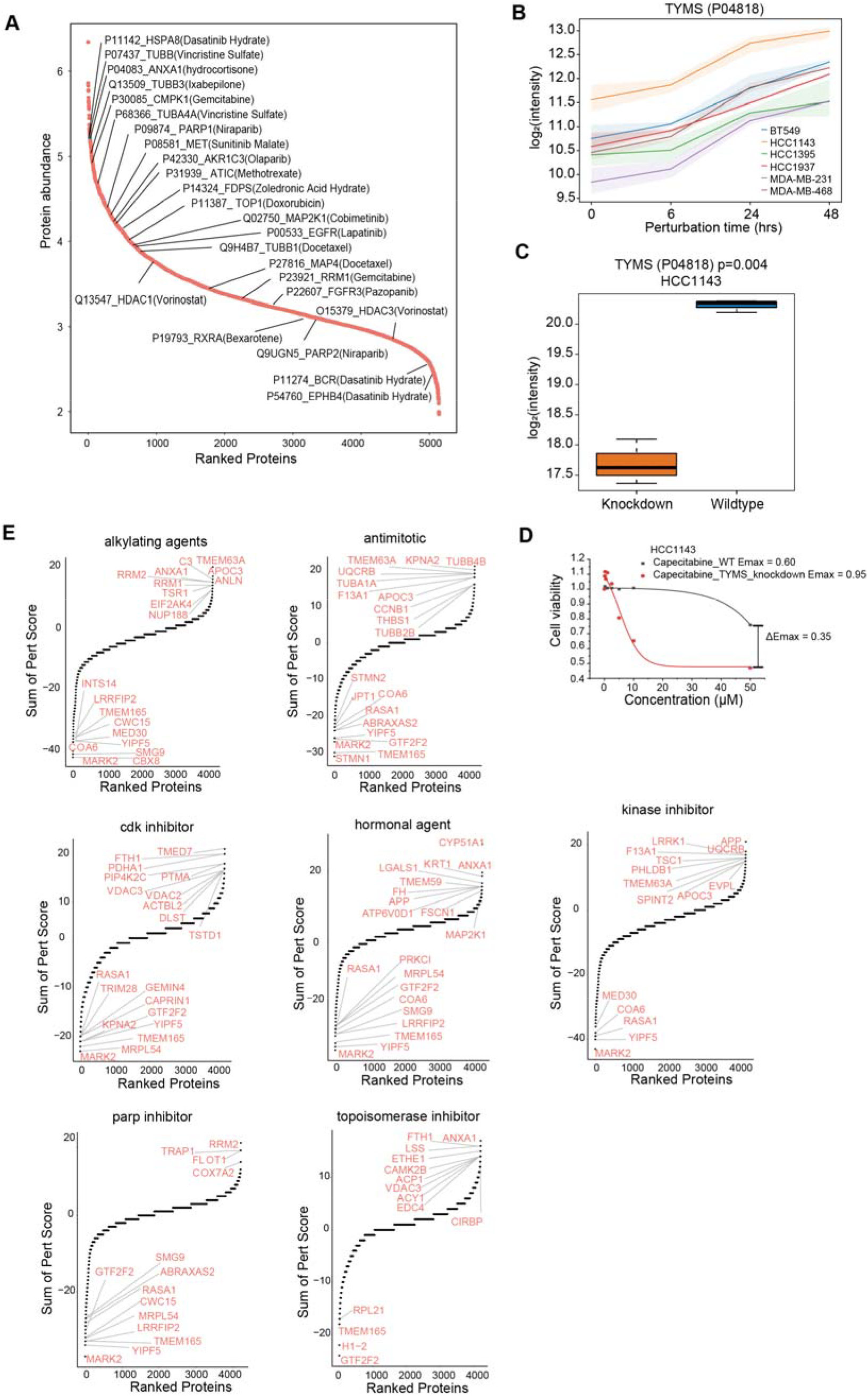
Dysregulated perturbation score. **(A)** The targeted proteins are identified in this perturbation proteomics dataset. **(B)** TYMS protein expression after treatment. **(C)** Box plots depicting the log2-scaled protein abundance detected by MS to evaluate RNA interference efficacy. The knockdown efficiency of TYMS was evaluated in HCC1143 cells. **(D)** Cytotoxicity assay results of HCC1143 cells interfered by TYMS siRNA then treated with capecitabine. **(E)** Perturbation score for different drug classes.

**Supplementary Figure 3.**
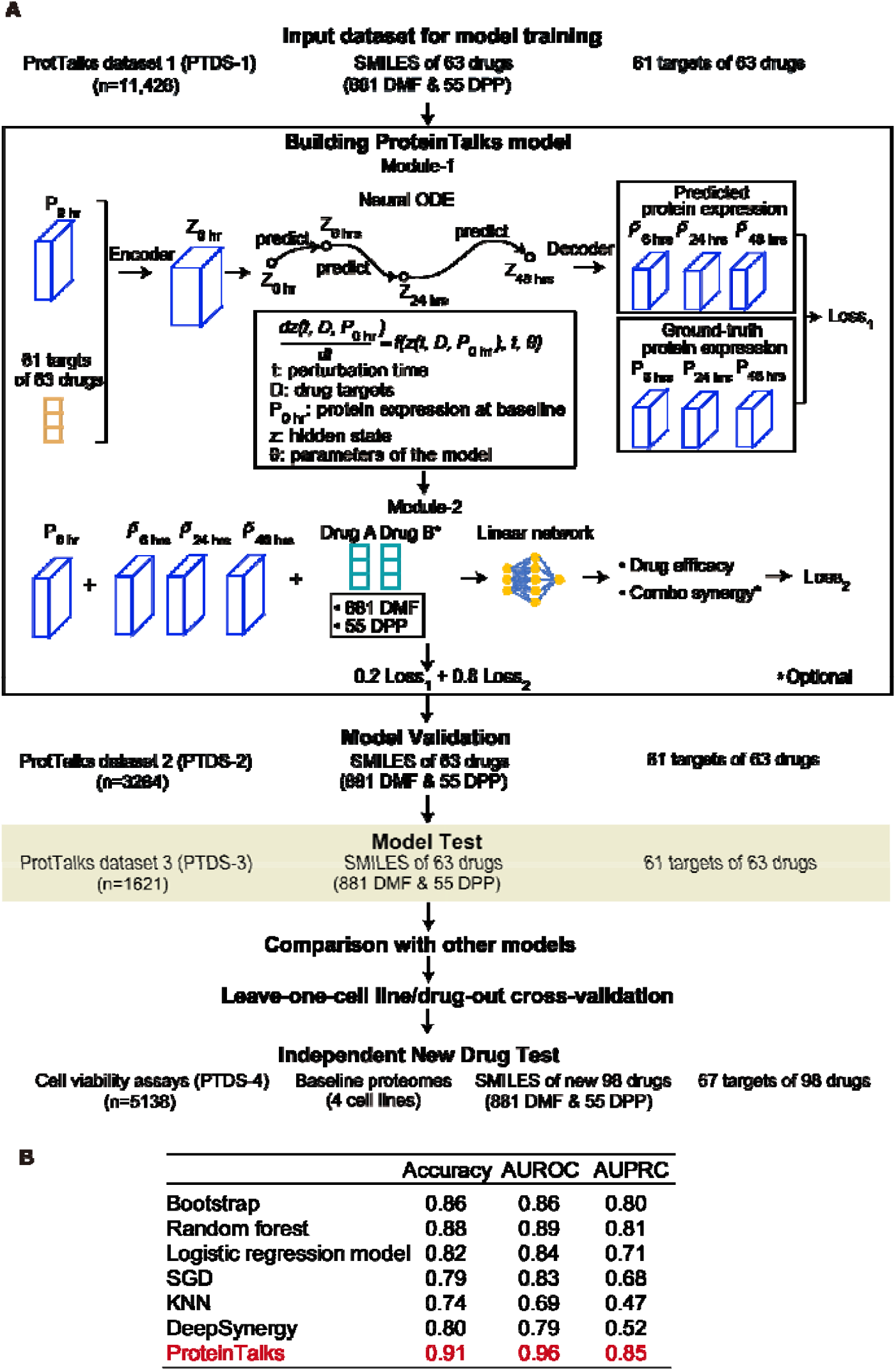
Detailed workflow of ProteinTalks building. Schematic representation of the process involved in constructing the ProteinTalks model. The 16,311 perturbation proteomic data were randomly divided into a training dataset (PTDS-1, n=11,426), a validation dataset (PTDS-2, n=3264), and a test dataset (PTDS-3, n=1621), following a 7 : 2 : 1 split. After data preprocessing, PTDS-1 was used for training the model through two modules, while PTDS-2 as validation set was used to optimize the model parameters. The performance of the ProteinTalks model was evaluated using PTDS-3 and compared with other models (**Figure 3B**). Leave-one-cell-line-out was used for cross-cell-line-validation (**Figure S4**), while leave-one-drug-out was used for cross-drug-validation (**Figure S5A-C**). Finally, a set of 98 new drugs was independently assayed for cell viability in four TNBC cell lines. This set was used to evaluate the predictive efficacy of the ProteinTalks model for new drugs (**Figure S5D**). DMF, dimensional drug molecular fingerprints; DPP, dimensional drug physicochemical properties.

**Supplementary Figure 4.**
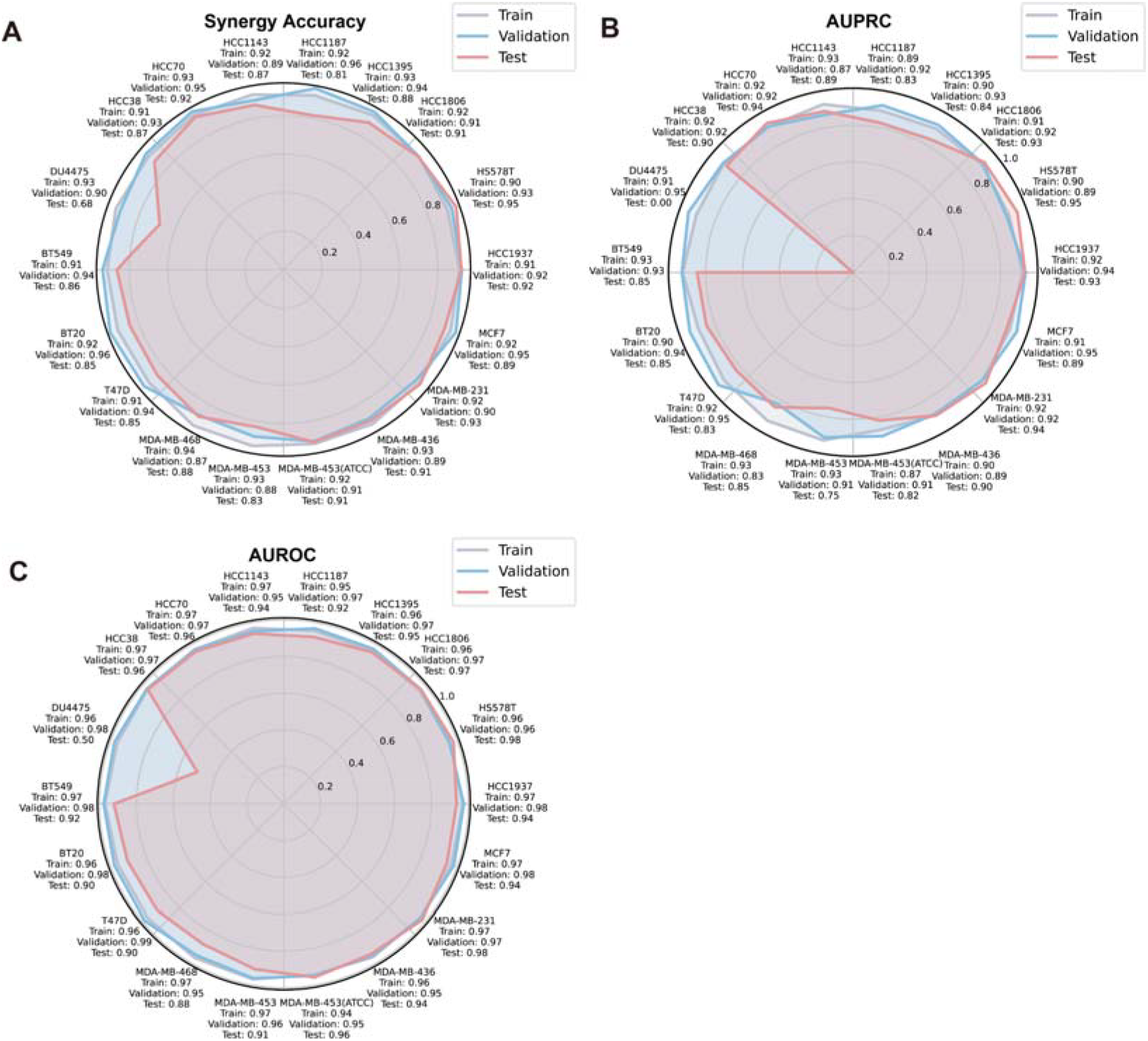
Leave-one-cell line-out cross-validation of ProteinTalks. **(A)-(C)** Accuracy **(A)**, AUPRC **(B)**, and AUROC **(C)**. The radial plots display the performance metrics for each cell line, including Synergy Accuracy, AUPRC (Area Under the Precision-Recall Curve), and AUROC (Area Under the Receiver Operating Characteristic Curve). Each plot shows the results for the training, validation, and test sets, indicated by gray, blue, and red lines, respectively. Performance metrics are presented for each cell line, indicating the model’s ability to generalize across different contexts and datasets. These metrics are crucial for assessing the predictive power and robustness of the ProteinTalks model in protein interaction and drug efficacy studies.

**Supplementary Figure 5.**
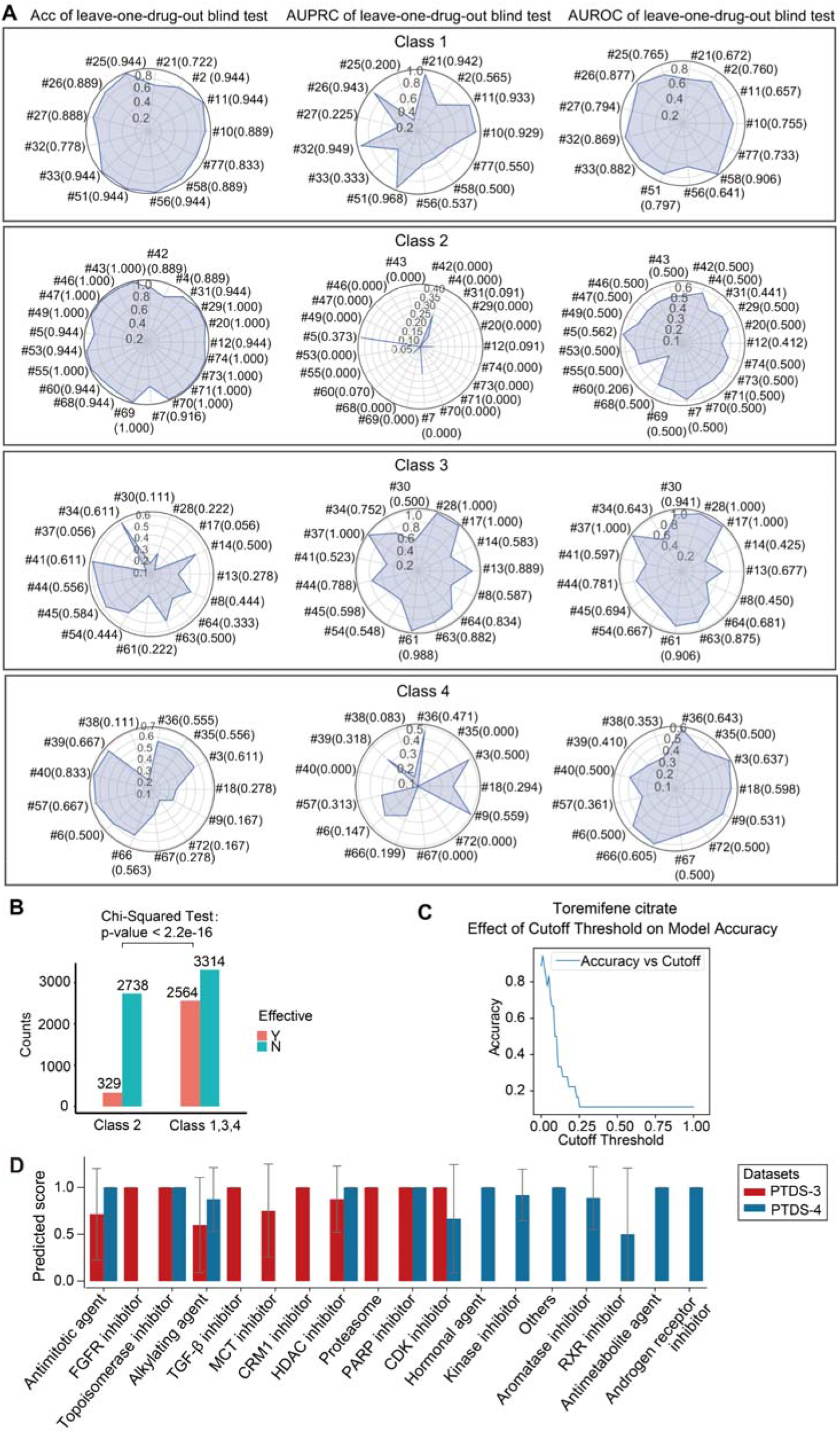
Testing of new drugs using ProteinTalks. **(A)** The four classes of drugs are defined by their accuracy (Acc), AUPRC, AUROC, which are determined through leave-one-drug-out cross-validation. The radial plots display the performance metrics for each cell line, including Synergy Accuracy, AUPRC (Area Under the Precision-Recall Curve), AUROC (Area Under the Receiver Operating Characteristic Curve), and Proteomics Multi-time Correlation. Each plot shows the results for the test datasets and the median value are shown in the brackets. Performance metrics are presented for each cell line, indicating the model’s ability to generalize across different contexts and datasets. The metrics are crucial for assessing the predictive power and robustness of the ProteinTalks model in protein interaction and drug efficacy studies. **(B)** The curve shows the effect of the cutoff threshold on the accuracy of the ProteinTalks model for predicting the efficacy of toremifene citrate. **(C)** Columns display the counts of effective drugs and ineffective drugs for each cell line in class 2 alone, as well as in combination with class 1, 3, 4. The Chi-Squared test was used to compare the differences between the effective and ineffective drugs for each cell line in two groups: class 2 alone, and the combination of class 1,3,4 (p-value < 2.2e-16). **(D)** Comparative display of prediction scores for each drug group generated by the ProteinTalks model for PDTS-3 and PDTS-4.

**Supplementary Figure 6.**
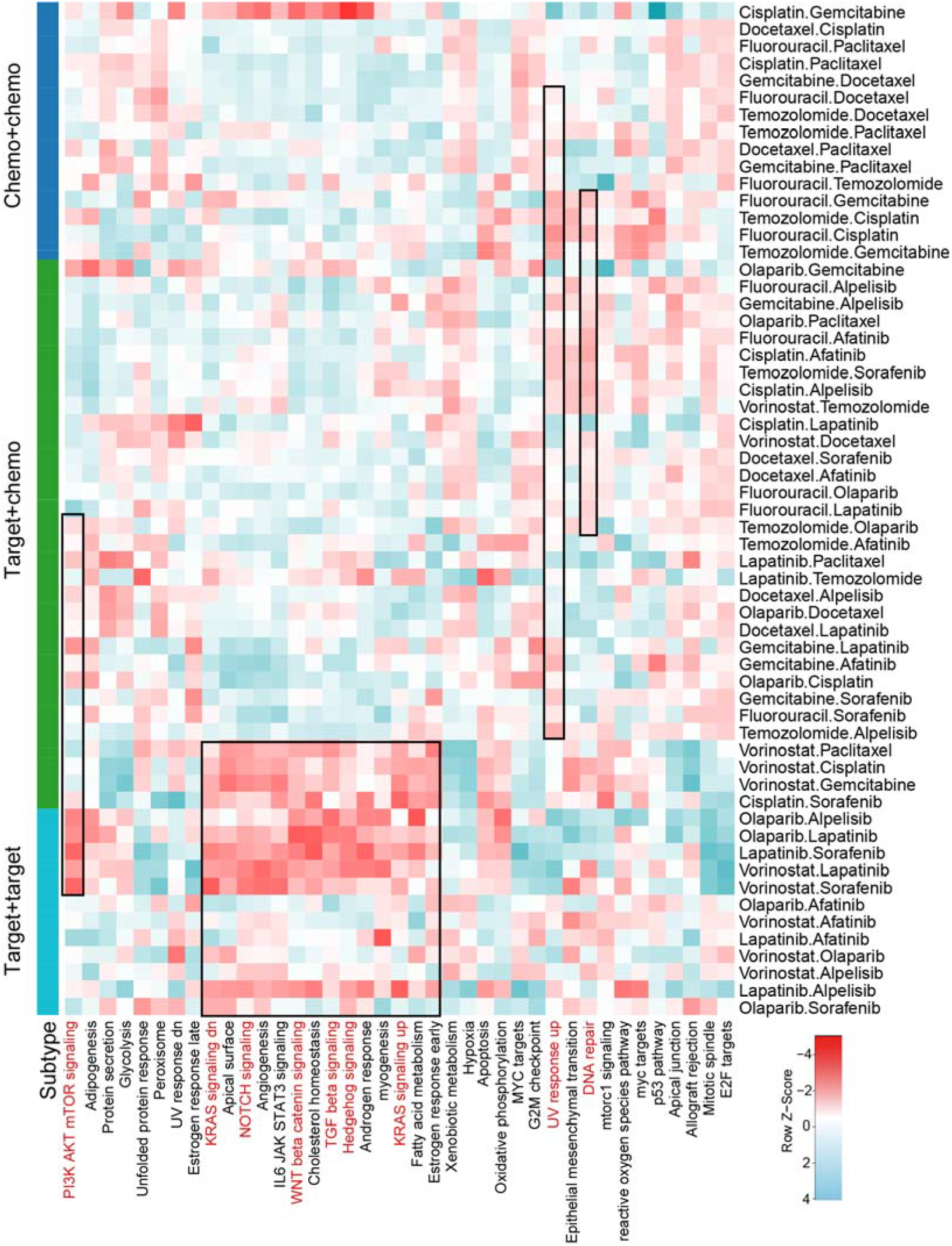
Pathway SHAP values for drug combinations. This heatmap shows the normalized pathway SHAP values, which is calculated using ProteinTalks for drug combinations after the training process. Heatmap shows the SHAP values of each pathway for each drug combination after z-score normalization. Each row represents different protein pathways, and each column represents different drug combinations. Hierarchical clustering was performed using Euclidean distance and the complete linkage agglomeration method. Different colors in the columns indicate different categories of drug combinations.

**Supplementary Figure 7.**
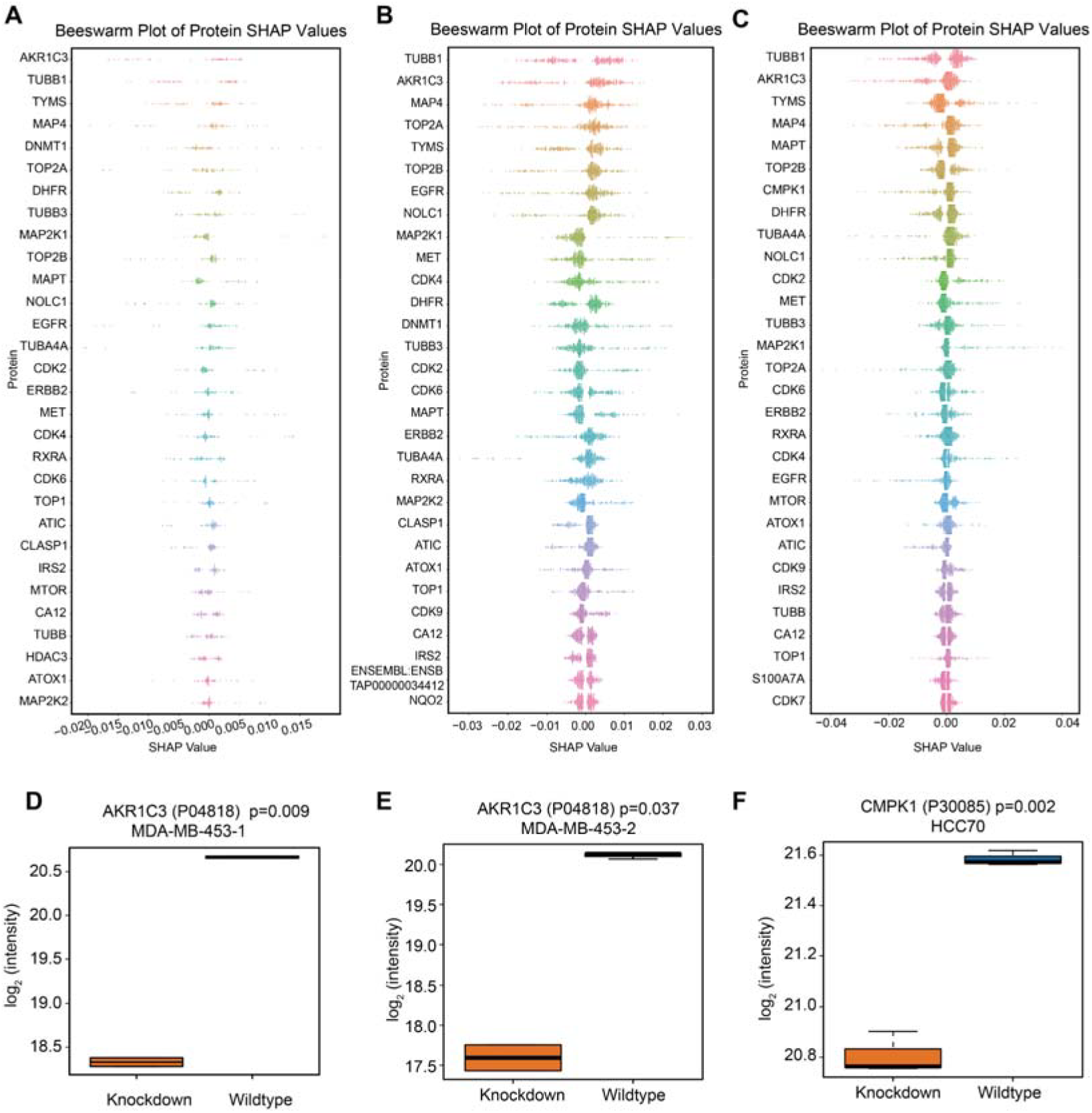
Interpretation of ProteinTalks model using SHAP values for different drugs. **(A)-(C)** Beeswarm plots illustrate the protein-level contributions to the model’s predictions for various drugs, including hormonal agents **(A)**, kinase inhibitors **(B)**, and alkylating agents **(C)**. Each point represents an individual protein’s contribution, with different colors indicating different proteins. The x-axis represents the SHAP value, while the y-axis displays the corresponding proteins. Positive SHAP values signify a positive contribution to the predicted efficacy, whereas negative values indicate a negative contribution. The absolute value of the SHAP score represents the magnitude of the contribution, with larger absolute values indicating a more significant influence on the predicted efficacy. **(D)-(F)** Box plots depicting the log2-scaled protein abundance detected by MS to evaluate RNA interference efficacy. The knockdown efficiency of AKR1C3 was evaluated in MDA-MB-453-1 cells **(D)**, MDA-MB-453-2 cells **(E)**, and CMPK1 in HCC70 cells **(F)**.

**Supplementary Figure 8.**
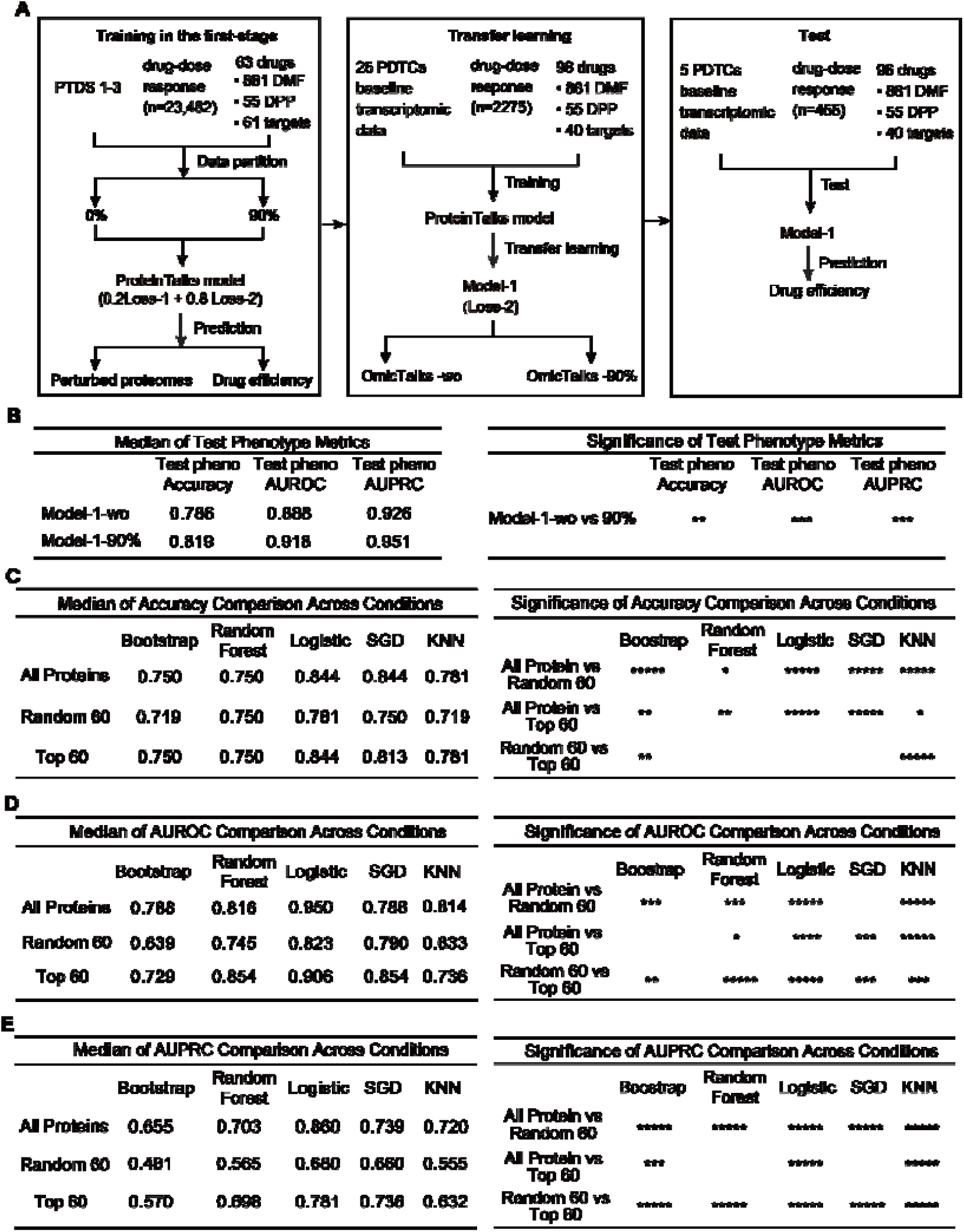
Assessing the clinical relevance of the model-1 with PDX transcriptomics data. **(A)** Diagram illustrating the construction of the ProteinTalks model. The model underwent training with perturbation proteomic data and was subsequently refined using a subset of transcriptomic data from PDX models. The remaining transcriptomic data were used to evaluate the model’s capability to predict drug efficacy. Model-1-wo, Model-1-without, was trained from scratch solely on the same subset of transcriptomic PDTC data without transfer learning from the perturbation proteomic data. When 90% of the PTDS-1, -2, and -3 perturbation proteomic data were used in the first stage, the corresponding models obtained were Model-1-wo and Model-1-90%, respectively. **(B)** Evaluation of the model’s drug efficacy prediction performance, with pretraining conducted using varying percentages of perturbation proteomic data. Statistical significance was determined via the Mann-Whitney U test. **(C)-(E)** Performance comparison of drug combination synergy predictions in TNBC patient cohorts, utilizing the complete proteome, a random selection of 60 proteins, and the top 60 proteins identified by SHAP values within the ProteinTalks. Significance levels were assessed using the Mann-Whitney U test, with p-values denoted as follows: *, p < 0.05; **, p < 0.01; ***, p < 0.001; ****, p < 0.0001.

**Supplementary Figure 9.**
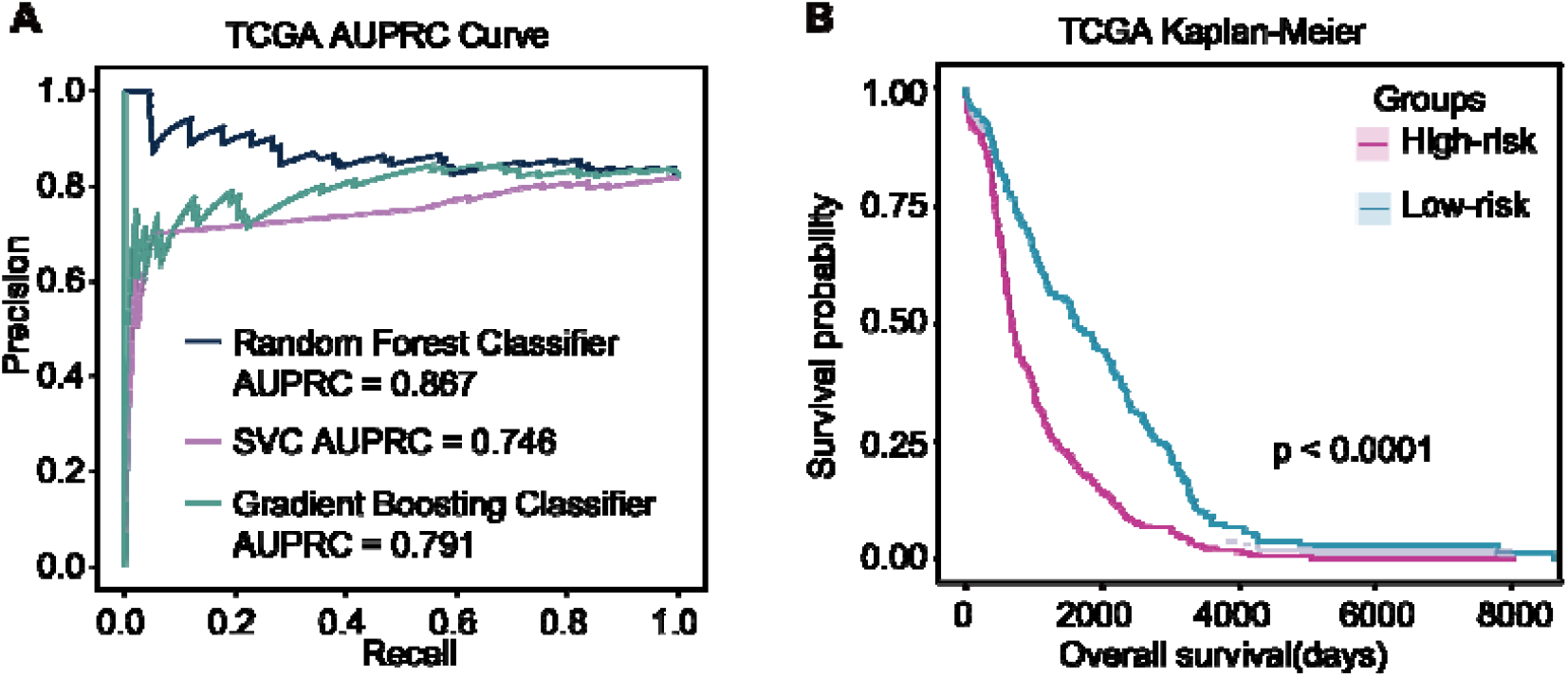
Clinical relevance validation in TCGA dataset. **(A)** In the TCGA transcriptomic dataset (N=823), the AUPRCs for the prognosis prediction performance in the test dataset (N=165) were evaluated using different machine learning models based on the top 60 proteins identified by ProteinTalks. **(B)** In the TCGA proteomic dataset, the Kaplan-Meier (KM) curves of overall survival based on the top 60 proteins identified by ProteinTalks.

